# *In vivo* multimodal lineage tracing of mammalian development by DeepTrack barcoding

**DOI:** 10.64898/2026.08.29.748052

**Authors:** Chenyu Guo, Junyao Jiang, Xiaomin Wang, Xinghuai Huang, Shenghu Zhang, Chenyang Shao, Mingyuan Zhang, Xiwen Hu, Wenqian Yang, Fuwei Shang, Xi Wang, Hongbo Zhai, Quan Du, Fang Liu, Danyang He, Xiaodong Liu, Guangdun Peng, Saifeng Cheng, Yanxiao Zhang, Duanqing Pei, Weike Pei

## Abstract

A comprehensive recording of cell fate transitions and underlying molecular changes remains a fundamental goal in developmental biology. Here, we present DeepTrack, a lineage tracing mouse model that integrates *in situ* cellular barcoding with high-throughput, single-cell multi-omics to simultaneously profile clonal fates, transcriptomic states, and chromatin accessibility. Using DeepTrack, we profiled clonal behaviors during gastrulation and early organogenesis, uncovered early fate priming within epiblast clones, and revealed clonal architecture within distinct regions of the nervous system. Embryo-wide multi-omic lineage tracing at single-cell resolution revealed transcriptional and epigenetic programs underlying fate commitment in neuromesodermal progenitors (NMPs). Clonal tracing with multi-omic profiles enabled inference of fate-associated gene-regulatory networks and identified the transcription factor *Cdx2* as a key regulator of mesodermal specification in NMPs. Genetic perturbation of *Cdx2* in chimeric embryos impaired paraxial mesoderm differentiation. Together, DeepTrack provides a versatile framework for decoding multimodal regulation of cell fate across diverse developmental contexts.

---

Mammalian embryogenesis coordinates cellular processes such as proliferation, differentiation, and fate specification to generate all tissues and specified cell types. During this process, the epiblast, a transient cell population in early mammalian development, gives rise to the three germ layers through gastrulation^1^. Following gastrulation, neurulation represents the next major developmental event, which proceeds via two distinct modes. Primary neurulation involves the folding and fusion of the neuroectoderm-derived neural plate to form the anterior neural tube^2^. In contrast, secondary neurulation is driven by NMPs located in the caudal region of the embryo that generate neural cells, thereby elongating the neural tube along the anterior-posterior axis^3–7^. The complete neural tube ultimately gives rise to the brain and spinal cord.

A comprehensive recording of changes in cell states, locations, and fate outcomes can reveal the order of cellular events and molecular mechanisms driving cell fate choice. Recent advances in single-cell transcriptomics and epigenomics enable high-resolution mapping of cellular differentiation dynamics by densely profiling cells across developmental stages^8–15^. While these technologies can be used to construct continuous trajectories that capture cell state transitions during differentiation, they may not directly link early cell states and terminal cell fates^16,17^. Therefore, state trajectories do not directly reveal how cells navigate between states or choose specific trajectories at developmental branch points^18^.

Lineage tracing has traditionally served as the gold standard approach to label individual cells at an early stage and track their clonal progeny over time^19^. Recently, lineage tracing methods have advanced to track cell clones using induced and heritable DNA tags (barcodes) analyzed through sequencing, enabling high-throughput, simultaneous tracking of multiple lineages *in vivo*^20–24^. By integrating barcoding and single-cell transcriptomics, single-cell lineage tracing enables the simultaneous capture of lineage information and molecular state in the same single cell^25–37^. The application of these approaches has revealed cell fate heterogeneity and identified novel regulators underlying cell fate choice^38^. Notably, changes in the epigenome, such as chromatin accessibility, are also critical for regulating gene expression and cell fate^39,40^. During differentiation, epigenetic shifts often precede changes in gene expression^41,42^. Gene regulatory networks (GRNs) reflect the complex interplay between transcriptome and epigenome, representing core regulatory programs that govern cell fate decisions. This underscores the need for tools that simultaneously resolve cell fate, transcriptomic, and epigenomic states in single cells. While recent studies have started to achieve this using single-cell multi-omics, these efforts have largely focused on *in vitro* models or tissue samples with limited cell numbers^43,44^. A major technical challenge remains the joint capture of lineage, transcriptomic, and epigenomic information for *in vivo* systems with large cell populations, such as developing embryos.

To address this challenge, we present a lineage tracing mouse model (DeepTrack), which enabled us to generate a comprehensive lineage-traced single-cell landscape of mammalian development. More importantly, DeepTrack is compatible with a high-throughput single-cell multi-omics platform, thereby enabling simultaneous profiling of lineage barcodes, chromatin accessibility, and gene expression from a large number of single cells *in vivo*. As a proof-of-concept, we applied DeepTrack to profile clonal behaviors in mouse embryos, revealing the restricted contributions of epiblast clones to specific embryonic lineages. We used single-cell fate map of early organogenesis to reconstruct clonal relationships within spatially distinct regions of the neural tube and revealed previously unrecognized lineage relationships and fate restrictions. Focusing on the trunk region, we found that neuromesodermal progenitors (NMPs) are highly heterogeneous, with most clones biased to either paraxial mesoderm or spinal cord. Lastly, the addition of epigenomic information to clonal tracing data allowed us to construct fate-associated GRNs and to identify *Cdx2* as a critical regulator of NMP fate decisions. Collectively, we demonstrated the ability of DeepTrack to resolve lineage hierarchies across tissue regions, quantify clonal fate contributions, and uncover the interplay between the transcriptome and epigenome in cell fate choice. Our work establishes a versatile platform for multimodal *in vivo* lineage recording, applicable to diverse biological contexts.

## Results

### Single-cell clonal tracing of gastrulation and early organogenesis

We previously developed PolyloxExpress, a Cre-dependent RNA barcoding system that resolves cell fates and single-cell transcriptomes *in situ*^33^. We hypothesized that increasing lineage barcode expression could improve barcode detection in single-cell assays, thereby providing a viable route to multi-omic lineage tracing at single-cell resolution^18^. To enhance barcode expression *in vivo*, we generated DeepTrack mice (*Rosa26^DeepTrack/+^*), in which the Polylox barcode substrate was expressed by the strong CAG promoter (Fig. 1a). As expected, DeepTrack mice exhibited significantly higher barcode expression levels as measured by real-time quantitative PCR (Extended Data Fig. 1a). To evaluate *in vivo* barcoding efficiency in DeepTrack mice, we crossed the *Rosa26^DeepTrack/+^* line with *Rosa26^CreERT2^* mice bearing ubiquitously expressed, tamoxifen-inducible Cre and administered 4-Hydroxytamoxifen (4-OHT) at embryonic day 9.5 (E9.5), which transiently activated Cre in embryonic cells (Extended Data Fig. 1b). Our results show that DeepTrack barcodes were generated within a few hours *in vivo* and stably maintained thereafter (Extended Data Fig. 1c,d).

**Fig. 1.**
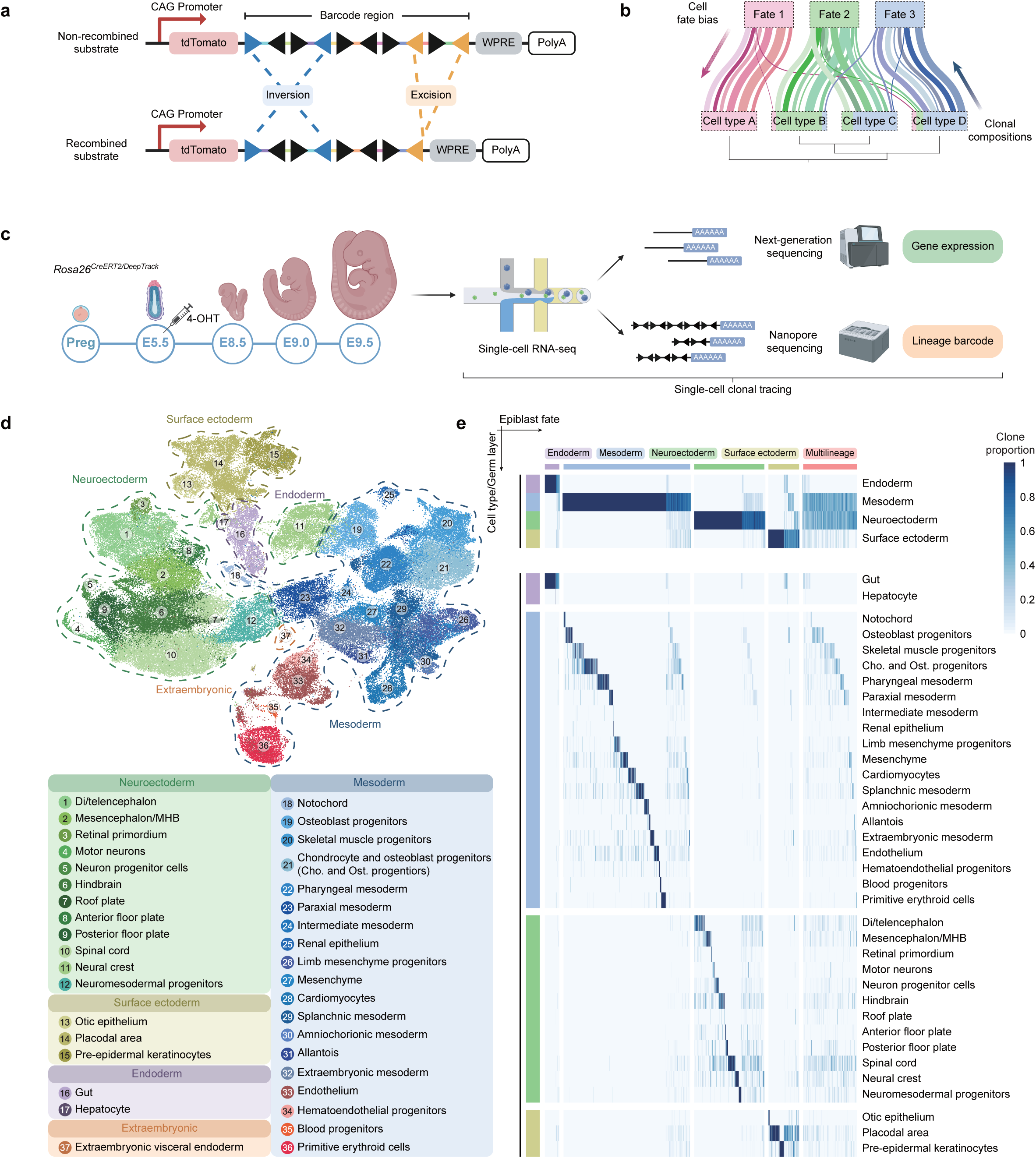
*In vivo* single-cell clonal tracing of gastrulation and early organogenesis by DeepTrack barcoding. **a**, Schematic of the DeepTrack barcoding system. The barcoding substrate contains 10 loxP sites (triangles) arranged in alternating orientations, flanking 9 unique DNA elements (colored linkers). Cre-mediated recombination results in random excisions (orange) and inversions (blue) to generate diverse genetic barcodes. Barcodes are transcribed as mRNAs under CAG promoter regulation. **b**, Framework for cell fate and lineage origin analysis, with lines represent individual clonal differentiation paths. **c**, Schematic of single-cell clonal tracing. Lineage barcodes were induced in *Rosa26^CreERT2/DeepTrack^*embryos before gastrulation. Whole embryos were harvested at indicated stages (E8.5, *n=*2; E9.0, *n=*1; E9.5, *n=*2) and analyzed via scRNA-seq to jointly capture transcriptomes and lineage barcodes. **d**, Uniform manifold approximation and projection (UMAP) displays 113,010 cells from single-cell clonal tracing data, color-coded by annotated cell types. **e**, Clone distributions across germ layers and cell types in five embryos. Columns represent individual clones, and rows denote germ layers or cell types. Clonal fate biases (top boxes) were defined by K-means clustering, with color intensity reflecting clone proportion per germ layer or cell type.

Next, we applied DeepTrack to profile the clonal behaviors in mammalian development. During gastrulation, epiblast cells give rise to the three germ layers (ectoderm, mesoderm, and endoderm) that ultimately orchestrate the development of the whole body plan ^1^. Despite its central role in embryogenesis, the clonal dynamics underlying lineage specification *in vivo* remain incompletely understood. To quantitatively resolve clonal fate specification during gastrulation and early organogenesis, we performed single-cell clonal lineage tracing experiments to simultaneously capture lineage barcodes and transcriptomes in individual embryonic cells. Barcodes were induced in pre-gastrulation stage (E5.5-E6.0) in *Rosa26^CreERT2/DeepTrack^* embryos using 4-OHT, and then assessed at E8.5, E9.0, and E9.5 using single-cell RNA sequencing (scRNA-seq) (Fig. 1b,c). Altogether, we annotated 37 major embryonic cell types from five embryos (total 113,010 cells) using established marker genes^10,13^ and captured 366-640 barcodes per embryo (Fig. 1d, Extended Data Fig. 2a,b, and Supplementary Table 1,2). Notably, we found 53.73%-67.14% of cells per embryo retained lineage barcodes, demonstrating the robust single-cell detection efficiency of DeepTrack barcodes (Extended Data Fig. 2c). Barcode detection rates varied across cell types, with the highest robustness observed in the nervous system (Extended Data Fig. 2d). Prior to further analysis, we filtered the total barcodes using a previously established generation probability (*Pgen*)^22,45^ threshold to retain rare barcodes that were more likely initially induced in single cells (see methods).

The propagation of clonal barcodes reflects the differentiation preference of epiblast cells into different germ layers and cell types (Fig. 1e). We identified heterogeneous clonal fate outcomes and defined them as: (1) multilineage clones, whose barcodes were detected across multiple germ layers, and (2) germ layer-specific/biased clones, whose barcodes were enriched in a single germ layer or a single cell type. The latter included endoderm-biased, mesoderm-biased, neuroectoderm-biased, and surface ectoderm-biased clones. Temporal recording of lineage outcomes at different stages indicated that this fate restriction was observed as early as the onset of organogenesis (E8.5) (Extended Data Fig. 3a). At the clonal level, fate contributions were highly variable between clones (Extended Data Fig. 3b-f). To further validate this heterogeneous clonal contribution in embryonic organs at later stages, we barcoded *Rosa26^CreERT2/DeepTrack^* embryos at the pre-gastrulation stage (E5.5) using 4-OHT and collected 16 embryonic and extraembryonic tissues from two E18.5 embryos (Extended Data Fig. 4a). The heterogeneous distribution of epiblast-derived barcodes across germ layers and tissue types was observed again (Extended Data Fig. 4b,c), supporting the concept that clonal fate bias was established before organ formation.

### Clone reconstruction across regions in the developing nervous system

A fundamental question in neurulation is whether spatially adjacent regions reflect a common developmental origin. To address this question, we investigated clonal architecture within the nervous system, focusing on spatial regionalization along the anterior-posterior and dorsal-ventral axes. Using five *Rosa26^CreERT2/DeepTrack^* embryos from single-cell clonal lineage tracing experiments, we projected 551 clones across neural tube compartments spanning retinal primordium, forebrain (diencephalon/telencephalon), midbrain (mesencephalon), hindbrain, spinal cord, and NMPs (Fig. 2a,b). We observed that 31.58% of clones exhibited single-region restriction, while 25.59% accumulated within adjacent bi-regional domains (Fig. 2c and Extended Data Fig. 5a). These results revealed that progenitors generate progeny cells with limited regional distribution in the developing nervous system. Notably, the spinal cord emerged as a hub sharing clones with its anterior (hindbrain, 60 clones) and posterior (NMPs, 42 clones) compartments, indicating its unique role in bridging anterior-posterior regionalization (Fig. 2c).

**Fig. 2.**
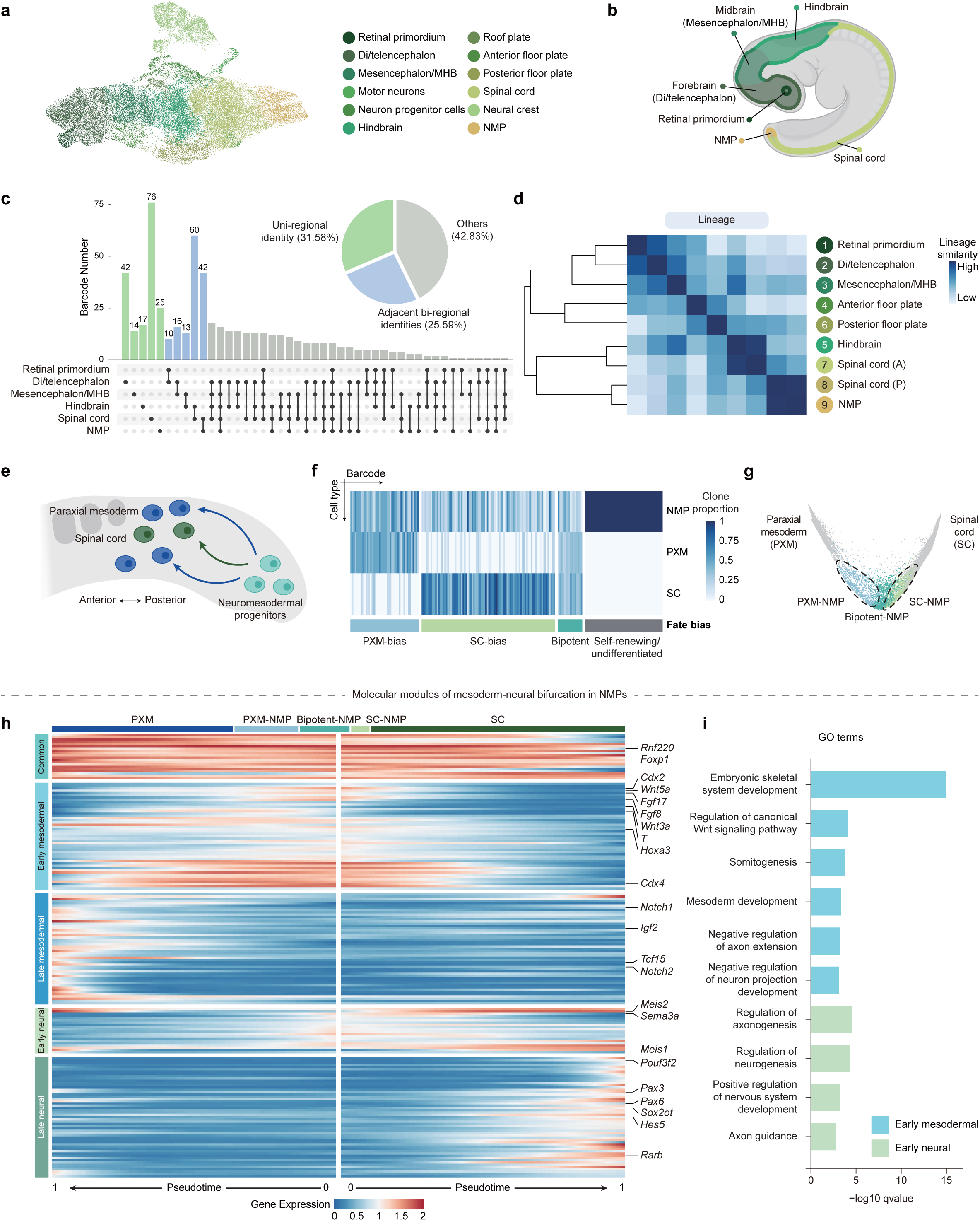
Clone reconstruction across the neural tube and clonal fate bias of NMPs in *vivo*. **a**, UMAP displays 46,087 neuroectoderm cells from single-cell clonal tracing data, color-coded by annotated cell types. **b**, Anatomical view of neural tube regions in an E9.5 embryo. **c**, Upset plots display clonal overlaps across regions in the nervous system. Clonal identities are color-coded as follows: uni-regional (green), adjacent bi-regional (blue), and other (gray) (*n=*5 embryos). **d**, Hierarchical clustering of cell types based on lineage barcode similarity (*n*=5 embryos). Color gradients indicate similarity scores among cell types. A, anterior; P, posterior. **e**, Schematic of embryonic trunk regions at the onset of organogenesis. **f**, Distribution of clonal barcodes (columns) from NMPs to PXM or SC lineages (rows). Only the 200 barcodes labeling NMPs are displayed. Clonal fate biases (bottom boxes) were defined by K-means clustering, with color intensity reflecting clone proportion per cell type. **g**, CoSpar-predicted fate bias for NMPs transitioning to mesodermal or neural lineages, classifying bipotent NMPs, PXM-NMPs, and SC-NMPs. **h**, Heatmap of gene expression dynamics along NMP differentiation trajectories. Rows represent differentially expressed genes (DEGs), and columns represent meta-cells ordered by pseudotime. Left boxes denote gene modules defined by K-means clustering, and top boxes indicate cell types. **i**, Gene Ontology (GO) enrichment of early mesodermal or neural modules, highlighting fate-associated biological pathways.

To map clones along the anterior-posterior axis, we performed integrated analysis of single-cell and spatial transcriptomics and resolved spinal cord subclusters, with spinal cord (A) mapped to the anterior side and spinal cord (P) aligned with the posterior side (Extended Data Fig. 5b,c). Next, we reconstructed a lineage tree of the nervous system based on barcode similarity (Fig. 2d). This lineage analysis for cells with positional identity revealed that the forebrain and midbrain shared a common lineage, whereas the hindbrain likely originated from a separate developmental pathway shared with spinal cord (A) (Fig. 2d). Clonal barcode similarity between hindbrain and spinal cord (A) was observed as early as E8.5, probably arising from common progenitors during the formation of anterior-posterior axis^23^ (Extended Data Fig. 5d). In addition, we also found that barcodes from spinal cord (P) and NMPs were more closely related (Fig. 2d). To resolve clonal relationships along the dorsal-ventral axis, we used established markers (dorsal: *Zic1* and *Pax3*; ventral: *Nkx6-1* and *Nkx2-9*) to annotate hindbrain and anterior spinal cord subclusters ^46^ (Extended Data Fig. 5e-i). Clonal barcode similarity revealed that cells from the dorsal side of each tissue region were more developmentally related to cells from the ventral side of the same region (Extended Data Fig. 5j).

### Clonal fate specification of NMPs in *vivo*

To elucidate early divergence of germ layers, we reconstructed lineage hierarchies of embryos collected at E8.5, E9.0, and E9.5 (Extended Data Fig. 6a). In contrast to the conventional model that the three germ layers assemble the major branchpoints in differentiation, our barcoding data revealed a close relationship between the neuroectoderm and mesoderm, supporting a common progenitor persisting through this stage ^37,47^ (Extended Data Fig. 6a).

NMPs are the common origin of both paraxial mesoderm and spinal cord in the trunk (Fig. 2e), which may partially explain the observed barcode similarity between the mesoderm and neuroectoderm ^6,48^. To quantitatively resolve the contribution of NMPs to mesodermal and neural lineages *in vivo*, we performed single-cell lineage barcode analysis to map fate outcomes of individual NMP clones using our lineage tracing datasets (Fig. 2f). We categorized NMPs into four fate groups through clustering analysis of lineage barcode distribution: (1) self-renewing/undifferentiated (barcodes detected only in NMPs), (2) bipotent (barcodes propagated to both neural and mesodermal lineages), (3) spinal cord-biased (SC-NMPs, barcodes enriched in spinal cord), and (4) paraxial mesoderm-biased (PXM-NMPs, barcodes enriched in paraxial mesoderm) (Fig. 2f). Our results revealed that NMP clones constitute a highly heterogeneous population with early fate priming in *vivo* (Fig. 2f and Extended Data Fig. 6b-e).

Next, we elucidated the transcriptomic features of fate-biased NMPs. We obtained 414 NMPs with clonal barcodes after *Pgen* filtering. To further infer cell fate across the entire NMP pool, we employed CoSpar^49^, a computational approach robust to barcode dropout, which integrates single-cell transcriptomes with lineage information to predict fate biases (Fig. 2g, Extended Data Fig. 6f, and see methods). CoSpar predicted that bipotent NMPs were closer to the root of the transcriptional landscape compared to fate-biased NMPs (Fig. 2g). We then reconstructed the fate trajectory (pseudotime) from NMPs to differentiated cells using the above CoSpar-predicted fate probabilities and identified 189 differentially expressed genes along these trajectories that may regulate NMP differentiation (Fig. 2h). These genes were organized into five fate-associated modules based on expression trends across the two trajectories (Fig. 2h and Supplementary Table 3). Genes in the early mesodermal module included key regulators of mesodermal fate such as *Wnt* (*Wnt3a* and *Wnt5a*) and *Fgf* signaling (*Fgf8* and *Fgf17*), as well as somitogenesis-related genes like *Hox* genes (Fig. 2h). To assess whether these gene modules reflect changes in differentiation potential, we examined the biological function of fate-associated modules by Gene Ontology (GO) enrichment analysis. The early mesodermal module involved genes in the negative regulation of neural development, whereas the early neural module included genes promoting neurogenesis, reflecting competing molecular programs underlying NMP fate choice^50,51^ (Fig. 2i). Collectively, our results highlight the value of DeepTrack in identifying distinct transcriptional modules that may govern NMP lineage differentiation.

### Embryo-wide high-throughput multi-omic lineage tracing at single-cell resolution

Epigenomic dynamics, such as changes in chromatin accessibility, are known to regulate lineage specification during embryogenesis^9,52^. Integrative profiling of lineage, transcriptome, and epigenome within the same cell could provide deeper mechanistic insights into cell fate regulation. To achieve this, we developed a single-cell multi-omic clonal tracing platform based on DeepTrack. This platform incorporates lineage barcode measurement within single-cell co-assays for RNA and ATAC (Assay for Transposase-Accessible Chromatin using sequencing) modalities. Briefly, cellular mRNAs and expressed lineage barcode transcripts are retrieved from the individual nuclei through a modified library preparation protocol (see methods), while accessible chromatin regions from the same nucleus are profiled via the standard workflow for single-cell multi-omic sequencing (Fig. 3a).

**Fig. 3.**
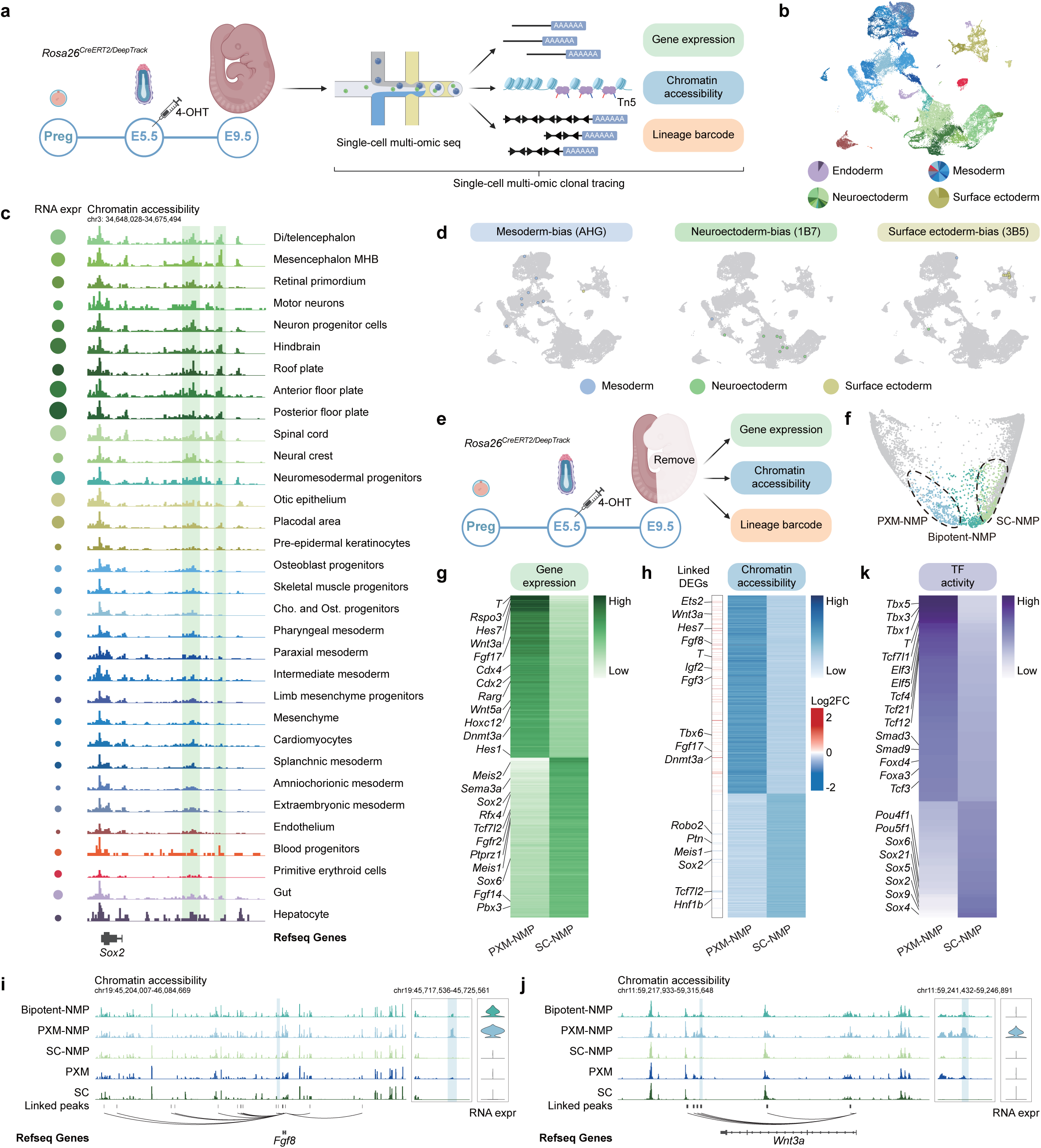
DeepTrack enables simultaneous *in vivo* recording of lineage, transcriptomes, and chromatin accessibility. **a**, Schematic of single-cell multi-omic clonal tracing. Lineage barcodes were induced in *Rosa26^CreERT2/DeepTrack^*embryos at E5.5. Two E9.5 embryos were profiled using single-cell multi-omics to capture gene expression, chromatin accessibility, and lineage barcodes. **b**, UMAP displays 60,355 cells from multi-omic lineage tracing data, color-coded by annotated germ layers and related cell types. **c**, RNA expression and chromatin accessibility of the *Sox2* locus across embryonic cell types. Each track shows pseudobulk ATAC-seq peaks for the indicated cell type. Dynamic distal regulatory regions (highlighted by green bars) exhibit cell-type specific accessibility patterns. Size of circle shows the expression level of *Sox2* (left). **d**, Representative clonal outcomes showing germ-layer biased clones in mesoderm, neuroectoderm, and surface ectoderm. **e**, Schematic of single-cell multi-omic clonal tracing for trunk tissues. **f**, CoSpar-predicted fate bias classifying 1,561 NMPs into bipotent, mesoderm-biased (PXM-NMPs), and spinal cord-biased (SC-NMPs) subtypes during trunk development. **g**, Heatmap of genes differentially enriched in PXM-NMPs versus SC-NMPs. **h**, Heatmap of differentially accessible chromatin regions (DARs) between PXM-NMPs and SC-NMPs. Peaks are annotated to their linked DEGs. The color of each linked DEGs corresponds to its log2 fold change in expression between PXM-NMPs and SC-NMPs. **i**,**j**, Chromatin accessibility and RNA expression of *Fgf8* and *Wnt3a*. Tracks display pseudobulk ATAC-seq signals for three NMP subtypes and their progeny cells (PXM and SC). DARs enriched in PXM-NMPs (blue bars) are highlighted in the middle plot. **k**, Heatmap showing differential TF activity in PXM-NMPs versus SC-NMPs.

To validate our lineage tracing approach across both RNA and ATAC modalities, we barcoded *Rosa26^CreERT2/DeepTrack^* embryos at E5.5 using 4-OHT treatment and profiled two E9.5 embryos using single-cell multi-omic sequencing (Fig. 3a). After quality control, we obtained 60,355 cells and identified all major lineages within both RNA and ATAC modalities (Fig. 3b, Extended Data Figs. 7,8 and Supplementary Table 4). We observed a mean of 6,172 UMIs (unique molecular identifiers) derived from 2,748 genes per cell, as well as 17,229 unique DNA fragments per cell, demonstrating high data quality across transcriptomic and epigenomic modalities (Extended Data Fig. 7a-c). By aggregating scATAC-seq data from multiple cell types into pseudobulk datasets, we confirmed that chromatin accessibility patterns of marker genes (e.g. *Sox2*, *Shh*, and *Prmt8*), particular at the distal intergenic and intronic regions, aligned with gene expression profiles (Fig. 3c and Extended Data Fig. 8c). Next, we performed barcode analysis on single-cell multi-omic clonal tracing data and identified 584 clones (Extended Data Fig. 9a). Consistent with our lineage tracing experiments, we reproduced the heterogeneous epiblast clonal fate pattern into distinct lineages and reconstructed a similar lineage tree (Fig. 3d, and Extended Data Fig. 9b,c).

To obtain more clones for mapping trunk development across RNA and ATAC modalities, we performed single-cell multi-omic sequencing on E9.5 trunk tissues from a *Rosa26^CreERT2/DeepTrack^* embryo barcoded at E5.5 (Fig. 3e). Integration of this trunk dataset with whole-embryo data enabled definition of PXM-NMPs and SC-NMPs based on lineage barcodes and CoSpar fate prediction (Fig. 3f and Extended Data Fig. 9d). We characterized fate-specific changes in gene expression and chromatin accessibility between these NMP fate groups. In the RNA modality, NMPs destined for mesodermal or neural fates exhibited significant transcriptomic differences (Fig. 3g). We identified 142 genes upregulated in PXM-NMPs, including the key regulator of mesoderm (*Brachyury*/*T*)^53^, the somite segmentation clock (*Hes7*) ^54^, and *Wnt* signaling (*Wnt3a* and *Rspo3*), as well as 140 upregulated genes in SC-NMPs, such as *Sox2* (Supplementary Table 5). In the ATAC modality, we observed distinct genome-wide chromatin conformation between PXM-NMPs and SC-NMPs, indicating extensive fate-specific epigenetic remodeling during cell fate determination (Fig. 3h and Supplementary Table 6). PXM-NMPs displayed 1,502 differentially accessible regions (DARs), whereas SC-NMPs exhibited 935 DARs. Fate-specific DARs showed significant enrichment in intronic and intergenic regions, suggesting that distal regulatory elements play a more prominent role in regulating cell fate decisions^43^ (Extended Data Fig. 9e,f). Next, we calculated the correlation between DARs and DEGs to identify putative fate-associated *cis*-regulatory elements (CREs). In PXM-NMPs, 75 upregulated DARs were linked to 43 upregulated DEGs, including mesodermal genes *Wnt3a* and *Tbx6* (Fig. 3h). SC-NMPs showed 20 enriched DARs correlated with 19 upregulated DEGs such as *Sox2* (Fig. 3h). As representative examples, *Fgf8* and *Wnt3a* showed differential expression in PXM-NMPs, accompanied by fate-specific chromatin accessibility increase at linked distal regulatory elements (Fig. 3i,j). Similarly, in SC-NMPs, *Sox2* and *Tanc2* were upregulated and exhibited fate-associated chromatin opening increase at intergenic regions (Extended Data Fig. 9g,h). These findings suggest that distinct CREs contribute to NMP fate choice by orchestrating fate-specific gene expression programs. To further uncover dynamic changes in transcription factor (TF) binding associated with fate-specific chromatin remodeling, we performed differential TF activity analysis to estimate the regulatory activity of TFs in different NMP fate groups. This revealed several lineage-specific TFs with increased activity scores, such as *T* in PXM-NMPs as well as *Sox2* in SC-NMPs (Fig. 3k and Supplementary Table 7). Together, these results indicate that fate-specific changes in gene expression are accompanied by extensive epigenomic remodeling.

### DeepTrack allows the inference of cell fate-associated GRNs and the identification of key TFs

Cell fate is governed by gene regulatory networks (GRNs) that represent the complex interactions between TFs, CREs, and target genes^55^. Fate-specifying TFs bind CREs (e.g. promoters and enhancers) to activate transcription of target genes essential for lineage commitment. Large-scale datasets with resolved lineage information and multimodal readouts empower reconstruction of fate-associated GRNs that capture core molecular programs directing cell fate choice^55^. Our high-throughput multi-omic lineage tracing approach thus provides a unique opportunity to decode fate-associated GRNs *in vivo*, systematically mapping TF-target regulations and identifying essential TFs for NMP fate specification (Fig. 4a). To achieve this, we constructed GRNs of PXM-NMPs and SC-NMPs using our multi-omic lineage tracing data^56,57^ (see methods). We assessed these fate-associated GRNs using degree centrality that measures the number of target genes directly connected to a specific TF. TFs with high centrality are likely essential for regulating molecular programs involved in fate specification^56^.

**Fig. 4.**
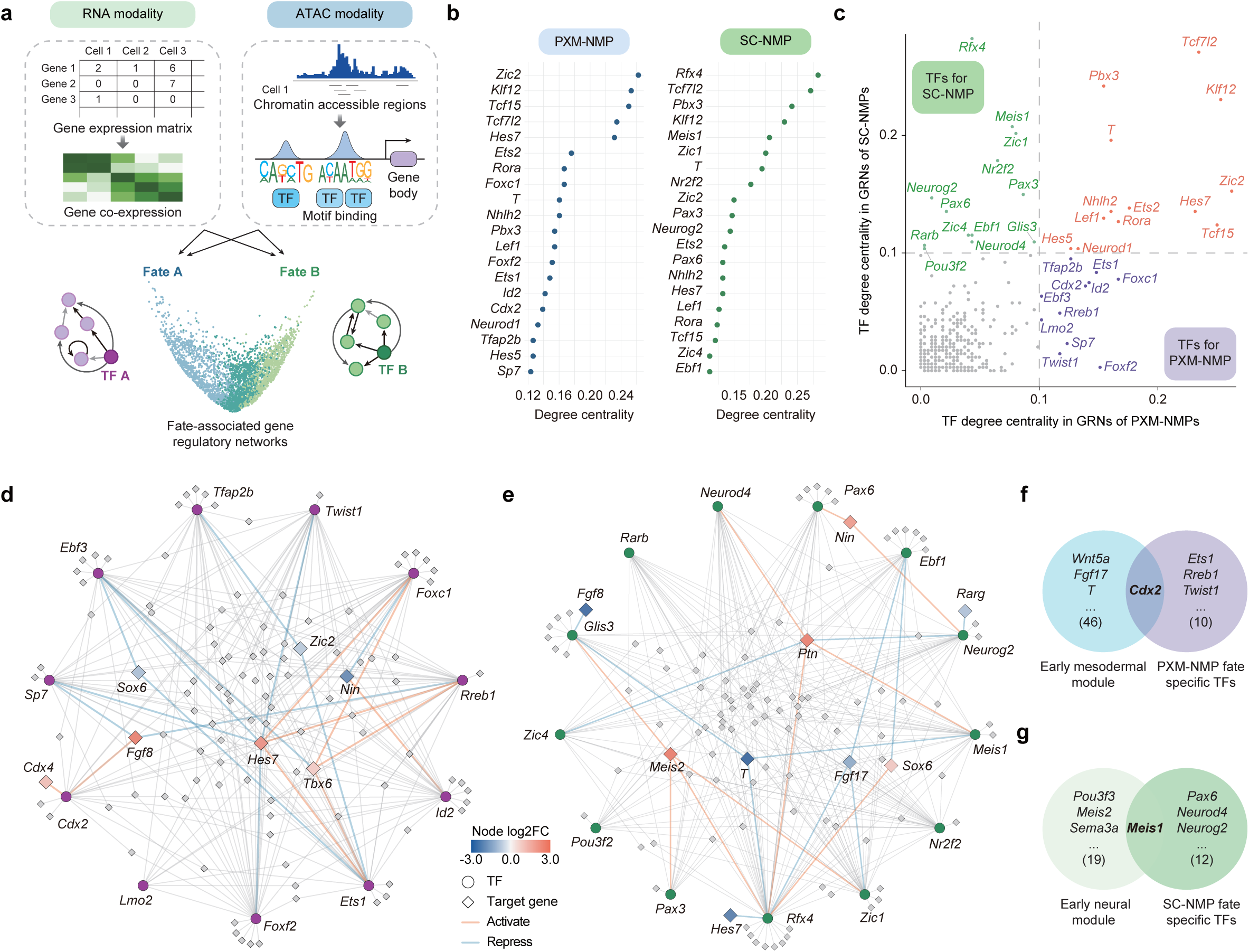
DeepTrack enables *in vivo* inference of fate-associated gene-regulatory networks and identification of key transcription factors. **a**, Workflow for the inference of fate-associated gene regulatory networks (GRNs). **b**, Degree centrality scores ranking top 20 transcription factors (TFs) in PXM-NMPs (left) and SC-NMPs (right). **c**, Scatter plot of key TFs regulating neural (green) or mesodermal (purple) differentiation in NMPs, with shared TFs denoted in red. **d**,**e**, Regulatory connections between fate-specific TFs and differentially expressed target genes in the GRNs of PXM-NMPs (**d**) and SC-NMPs (**e**). Each circle corresponds to a TF and each diamond represents a target gene. Selected target genes are highlighted and color-coded by expression level. Red edges denote activatory relationships (positive TF-target correlation), whereas blue edges represent repressive relationships (negative correlation). **f**,**g**, Venn diagrams showing overlaps between fate-associated gene modules and fate-specific TFs in GRNs for PXM-NMPs (**f**) or SC-NMPs (**g**).

We ranked the top 20 TFs in each fate-associated GRNs based on degree centrality scores, reflecting TF potential to regulate lineage-specific differentiation (Fig. 4b). These results were summarized on a scatter plot, revealing known and putative regulators of NMP fate choice (Fig. 4c). For example, TFs associated with neural lineage differentiation, such as *Pou3f2*, were enriched on the top left side of the plot (Fig. 4c). In contrast, TFs involved in mesodermal differentiation, including *Rreb1*, were located at the bottom right (Fig. 4c). We further visualized the interactions between fate-specific TFs and differentially expressed target genes within each fate-associated GRNs (Fig. 4d,e). In the GRNs of PXM-NMPs, mesodermal genes (e.g. *Hes7* and *Fgf8*) were activated by TFs such as *Rreb1* and *Cdx2* (Fig. 4d). In contrast, these two mesodermal genes were repressed by other fate-specific TFs in the GRNs of SC-NMPs (Fig. 4e). To prioritize candidate TFs for experimental validation, we overlapped the fate-specific TFs identified in fate-associated GRNs (Fig. 4c) with genes from fate-associated modules generated by single-cell clonal tracing (Fig. 2h). The resulting list highlighted *Cdx2* as an essential regulator of mesodermal fate, while *Meis1* emerged as a potential key regulator of neural fate (Fig. 4f,g). Overall, our results establish DeepTrack as a robust single-cell multi-omic clonal tracing platform and demonstrate its application in the identification of TFs regulating cell fate specification.

### *In vivo* analysis of *Cdx2* in trunk development and neuromesodermal progenitors

Validating cell fate regulators requires perturbation experiments. However, disrupting key developmental pathways, even conditionally, often causes severe tissue disorganization and confounding indirect effects. By contrast, mosaic perturbation through chimeric embryos minimizes such systemic distortions, enabling the functional study of genes with strong in *vivo* phenotypes^10^.

Homozygous *Cdx2*-mutant mice exhibit embryonic lethality between E3.5-E5.5, preventing functional analysis at later embryonic stages *in vivo*^58^. To circumvent this challenge, we generated chimeric mouse embryos by injecting tdTomato-labeled, *Cdx2*-mutant mouse embryonic stem (ES) cells into wild-type blastocysts (Fig. 5a). In these chimeras, host cells contributed to all embryonic lineages, enabling us to assess the specific effects of *Cdx2* deficiency in relatively healthy embryos (Extended Data Fig. 10a). Compared with control chimeras, *Cdx2*-mutant chimeras showed reduced contribution of tdTomato^+^ cells to the posterior end of the embryo, indicating an essential role of *Cdx2* in trunk development (Fig. 5b). To determine whether *Cdx2* deficiency caused lineage-specific developmental abnormalities, we sorted tdTomato^+^ (*Cdx2*-mutant) and tdTomato^−^ (wild-type) cells from three E10.5 *Cdx2*-mutant chimeras (22.1%-57.3% chimeric efficiency) for scRNA-seq (Fig. 5a and Extended Data Fig. 10b,c). We recovered 11,098 *Cdx2*-mutant cells that were distributed across almost all lineages compared to wild-type cells (Fig. 5c). We then compared the relative contributions of wild-type and *Cdx2*-mutant cells to each lineage. Comparisons between wild-type and *Cdx2*-mutant cells revealed that loss of *Cdx2* disrupted paraxial mesoderm formation but still maintained spinal cord development (Fig. 5d). Interestingly, *Cdx2*-mutant tdTomato^+^ cells produced fewer blood progenitors and primitive erythroid cells, likely due to deficient yolk sac circulation^59^ (Extended Data Fig. 10d).

**Fig. 5.**
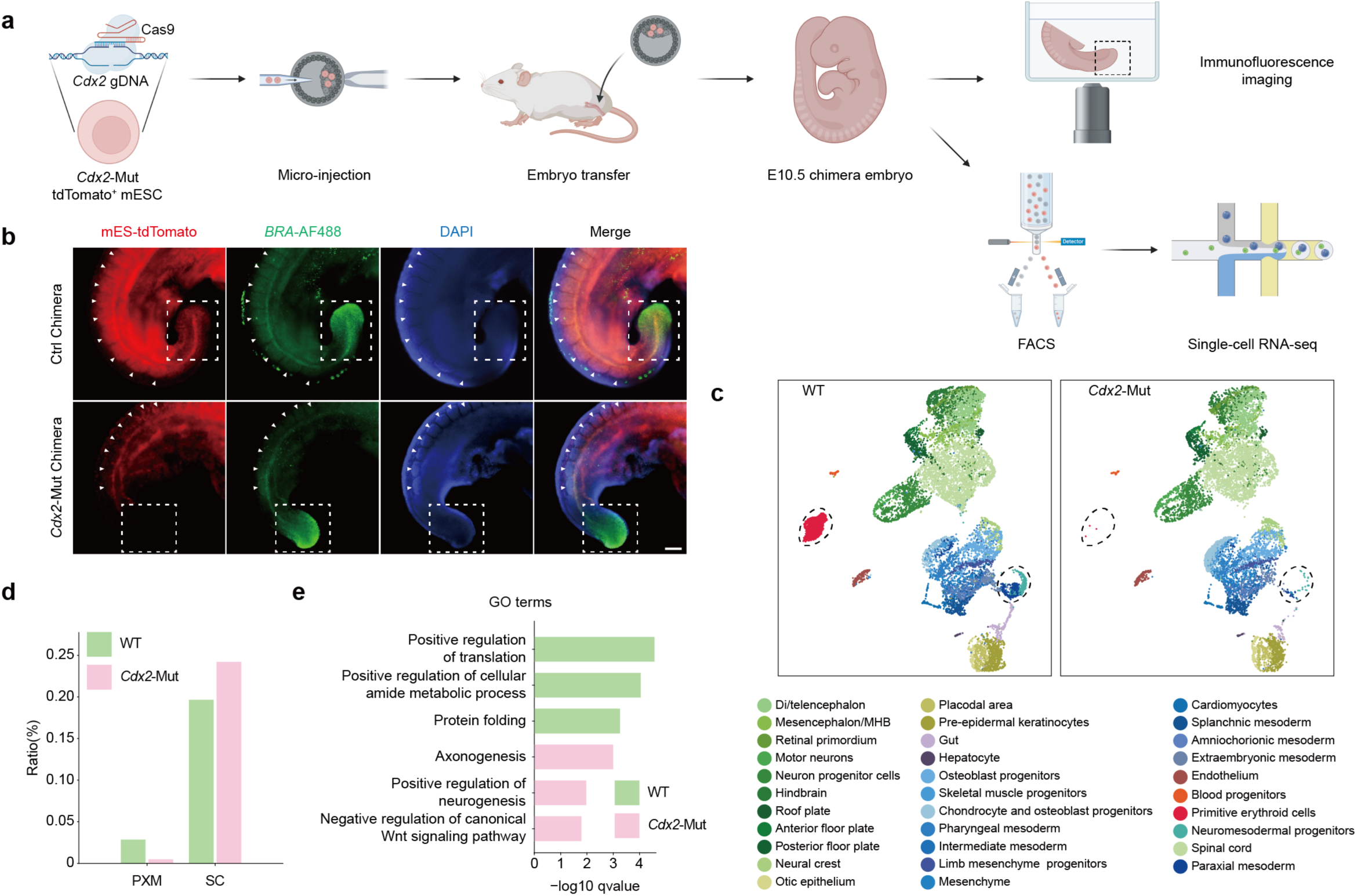
*In vivo* functional validation of *Cdx2* during development. **a**, Schematic workflow for generating *Cdx2*-mutant chimeras, followed by imaging and scRNA-seq. **b**, Immunofluorescence imaging of Brachyury/T in control (*n*=3) and *Cdx2*-mutant chimeras (*n*=3). Scale bar, 200 μm. **c**, UMAP displays wild-type (13,901 cells) and *Cdx2*-mutant (11,098) chimera cells, color-coded by annotated cell types. Dashed circles highlight major lineages disrupted in mutants. **d**, Quantified contributions of wild-type and *Cdx2*-mutant cells to paraxial mesoderm and spinal cord lineages. **e**, GO enrichment analysis of differentially expressed genes between wild-type NMPs and *Cdx2*-mutant NMPs.

To characterize transcriptional changes in *Cdx2*-mutant cells, we performed GO enrichment analysis and found that *Cdx2*-mutant NMPs upregulated genes promoting neurogenesis and inhibiting *Wnt* signaling (Fig. 5e and Supplementary Table 8). These results indicate that *Cdx2*-mutant NMPs activate a neural differentiation program. To further elucidate how *Cdx2* regulates NMP functions, we extracted potential *Cdx2* target genes from NMP fate-associated GRNs (Extended Data Fig. 10e). GO enrichment analysis for *Cdx2* target genes revealed that *Cdx2* represses neurogenesis-related genes, while activates mesodermal genes (Extended Data Fig. 10f). Together, these *in vivo* findings indicate a dual role of *Cdx2* as an activator of mesodermal genes and a repressor of neural genes within NMPs.

## Discussion

Resolving the cellular origins and molecular mechanisms of mammalian organogenesis is a central goal in both developmental biology and regenerative medicine. To address this, we developed DeepTrack, an integrative tool for high-throughput, multimodal clonal analysis *in vivo*. DeepTrack achieves highly efficient lineage barcode capture in single cells via traditional scRNA-seq and robust clonal fate mapping using single-cell co-assays for RNA and ATAC. Additionally, DeepTrack is compatible with inducible Cre lines, allowing temporally and spatially specific cellular labeling and lineage tracking across diverse tissues.

Using DeepTrack, we generated a comprehensive clonal atlas across all major tissues and cell types during early organogenesis and illustrated clonal fate diversity within epiblast cells over time. Contrary to the classic pluripotent model, where epiblasts contribute to all tissue types, our results revealed that substantial epiblast clones exhibit restricted contributions to specific germ layers or tissues *in vivo*. This phenomenon may arise from different timing of lineage-specific differentiation or spatial location divergence within epiblast clones^1,24^. As indicated by grafting experiments, epiblasts remain plastic at an early stage and then become fate restricted after they move through the primitive streak^60^. Clonal analysis indicated that epiblast position relative to the primitive streak serves as a robust predictor of their eventual fate^47,61,62^. Furthermore, epiblast cells from various regions could be transplanted to distinct locations and would acquire fates consistent with the adjacent host cells^60^.

Due to the technical difficulty of achieving large-scale in *situ* clonal tracing across the entire embryo, most high-throughput tracing efforts have predominantly been confined to specific tissue regions^34–36^. Our whole-embryo, high-throughput clonal tracking resolved a single-cell fate map of early organogenesis, providing a valuable resource broadly applicable to questions of tissue formation and its underlying clonal mechanisms. We exploited this resource by focusing on two key developmental events: the emergence of spatially divergent regions in the nervous system and the cell fate choice of NMPs in the trunk. We observed substantial clones accumulated in single or adjacent regions, suggesting that progenitors acquire positional identity before cell identity in the nervous system^63^. Further lineage analysis revealed separate lineage pathways between fore-midbrain and hindbrain, consistent with the lineage hierarchy reconstructed by CRISPR barcoding^23^. Cells in the hindbrain share greater clonal similarity with the anterior spinal cord than with the forebrain or midbrain, indicating that the hindbrain region is also derived from NMPs during secondary neurulation^64^. It remains to be tested if cells (fore-midbrain) with anterior identity are generated earlier, followed by cells (hindbrain and spinal cord) that emerge sequentially from a posterior region. Importantly, large-scale clonal analysis has uncovered hidden aspects of lineage relationships and fate restrictions. For instance, our study provides *in vivo* evidence that a spectrum of NMP fates coexists in the tail region, including self-renewing, bipotent, and lineage-biased populations toward spinal cord or paraxial mesoderm. In addition, our results revealed that mesodermal or neural lineage specification originates from transcriptionally distinct NMP populations, driven by competing molecular programs. NMP fate heterogeneity might be established by differential usage of developmental signals, such as *Wnt* and retinoic acid, when NMPs are exposed to distinct signaling environments at different positions^50,51^. In line with this, spatial transcriptomics of a human embryo shows that NMPs are organized into a bi-layered structure along the dorsal-ventral axis, with NMPs for mesoderm mainly on the ventral side, whereas NMPs for neuron are localized on the dorsal side of the NMP pool^64^.

Cell fate bias arises from both transcriptional and epigenomic heterogeneity, yet the molecular logic underlying this variation remain unclear. Identifying key factors of such biases is essential for uncovering the genetic and epigenetic mechanisms that orchestrate cell fate decisions in *vivo*. Here, we leveraged DeepTrack to simultaneously profile lineage barcodes, chromatin accessibility, and gene expression in single cells. This approach introduces an epigenomic dimension to complement current lineage tracing methods, establishing a foundation to find fate-specific regulatory elements and reconstruct fate-associated GRNs using clonal information. Focusing on NMPs, we found clear transcriptomic and epigenomic differences associated with fate outcomes. Through inference of fate-associated GRNs, we systematically assessed TF-target regulations and thus pinpointed *Cdx2* and *Meis1* as key TFs of PXM-NMPs and SC-NMPs, respectively. Several studies indicated the importance of *Cdx2* in trunk development^50,57,65,66^. Genetic perturbation of *Cdx* genes causes posterior body truncations *in vivo*^65,66^. *In vitro* culture systems, loss of *Cdx1/2/4* function disrupts posterior *Hox* gene expression, impairing mesoderm differentiation while promoting neural fate in NMP-derived tissues^50^. Our experiments using chimeric embryos further establish *Cdx2* as a critical regulator of trunk development and NMP function *in vivo*. Given the versatile role of *Cdx2* across multiple cell lineages, we cannot exclude the possibility that *Cdx2*-mutant cells might impair NMP-independent pathways or induce systemic developmental defects at early stages, which can also disrupt paraxial mesoderm development^59,63,65–68^. To address this, future studies could employ conditional perturbation strategies to achieve stage-specific *Cdx2* ablation *in vivo*.

Collectively, DeepTrack offers a robust and versatile platform for investigating diverse biological processes through multimodal lens. From a translational perspective, this approach could help uncover fundamental principles of clonal compensation in developmental disorders and regeneration, potentially illuminating the clonal basis of embryonic resilience. Despite the potential, our approach can be further improved in barcoding diversity and clonal recovery efficiency in single-cell assays. This might be achieved by generating multiplexed barcoding substrates at different genomic loci, as validated in CRISPR barcoding mouse models such as MARC1^23^, DARLIN^44^, and CREST^36^. In addition, current DeepTrack barcode size (400 bp - 2 kb) requires long-read sequencing, which remains less cost-effective for large-scale applications. Developing compact barcode substrates compatible with short-read platforms would be desirable^24^. While our study reveals fate diversity in epiblast cells and NMPs, it is not yet clear whether this heterogeneity is established by cell-intrinsic properties at the time of labeling or emerges from external cues. During embryogenesis, context-dependent signals orchestrate cell fate decisions to ensure the proper formation of tissues. Future technical advancements could integrate cellular barcoding with spatial transcriptomics and environmental sensors designed to detect cell-cell contacts or signaling history, facilitating the capture of interactions between clonal dynamics and microenvironmental changes within tissues^69–71^.

## Methods

### Mice

All animal procedures were approved by the Westlake University Institutional Animal Care and Use Committee (IACUC). *Rosa26^CreERT2^* mice^72^ (*B6.129-Gt(ROSA)26Sor^tm1(cre/ERT2)Tyj^/J*) were purchased from Jackson Lab (catalog JAX 008463). *Rosa26^PolyloxExpress^* (*B6-Gt(ROSA)26Sor^tm2.1(PolyloxExpress)Hrr^*) mice were gifted by Hans-Reimer Rodewald at the German Cancer Research Center (DKFZ, Heidelberg). Wild-type ICR mice were purchased from Zhejiang Vital River Laboratory Animal Technology Co., Ltd. The *Rosa26^DeepTrack^* mouse line was generated in this study. All primers for genotyping are provided in Supplementary Table 9.

### Generation of *Rosa26^DeepTrack/+^* knock-in mice

The *Rosa26^DeepTrack/+^* knock-in strain was generated by Shanghai Biomodel Organism Co., Ltd. In brief, the DeepTrack barcoding cassette was inserted into the *Rosa26* locus via CRISPR/Cas9-mediated homologous recombination. Cas9 mRNA and sgRNA were obtained through *in vitro* transcription, and the homologous recombination vector (donor vector) was constructed using In-Fusion cloning. This vector contains a 3.3 kb 5’ homologous arm, CAG-tdTomato-P2A-Polylox-WPRE-PolyA, and a 3.3 kb 3’ homologous arm. Cas9 mRNA, sgRNA, and the donor vector were microinjected into C57BL/6J zygotes to generate F0 founders. PCR amplification and Sanger sequencing were performed to identify positive F0 mice that were backcrossed with C57BL/6J mice to establish the F1 line.

### Barcode induction *in vivo*

For in *vivo* barcoding experiments, *Rosa26^DeepTrack/CreERT2^* embryos were generated by crossing *Rosa26^DeepTrack/+^* males with *Rosa26^CreERT2/CreERT2^* female mice. At E5.5 (5 days after the day of the plug), pregnant mice received a single dose of 4-OHT (Sigma, H6278; 0.4mg/mouse or 0.07 mg/g body weight) by oral gavage for barcode induction, together with 1.25 mg of progesterone (Sigma, V900699) to maintain pregnancy.

### Detection of lineage barcodes and transcriptomes in single cells

In single-cell clonal tracing experiments, *Rosa26^CreERT2/DeepTrack^* embryos were barcoded pre-gastrulation. Single-cell profiling included whole-embryos (E8.5 and E9.0) and partial E9.5 embryos. For all embryos, extra-embryonic tissues were removed, and somites were counted before embryo digestion. For E9.5 embryos, the head and the body were dissected from the branchial arch and processed separately. Dissected tissues were enzymatically digested with TrypLE Express (Thermo, 12605010) and filtered (40 µm) to generate single-cell suspensions in PBS with 0.04% BSA (Sigma, A1933) for scRNA-seq.

Lineage barcodes and transcriptomes were captured using 10X Genomics Chromium Next GEM Single Cell 3’ Reagent Kits v3.1, with modified protocols for barcode enrichment. In brief, a spike-in oligo (#ISPCR2999), annealing to the lineage barcoding cassette, was added to cDNA elution pre-amplification. In addition, the cDNA amplification program was adjusted to 3 min at 98°C; (15 s at 98°C, 20 s at 63°C, 3 min at 72°C) with 13-14 cycles; 3 minutes at 72°C.

For lineage barcode library preparation, targeted amplification of barcodes from a 10x cDNA library was performed via nested PCR using Expand Long PCR System (Roche, 11759060001). In the first round, we used primers #2,674 and #2999 for 3 min at 95 C; (30s at 95°C, 30 s at 60°C, 3 min at 72°C) 12 cycles; 10 min at 72°C. PCR products were purified with 0.7X SPRI beads (Beckman Coulter, B23318), and used for the second round of PCR in which we used primers #2,426 and #2,676 for 3 min at 95°C; (30 s at 95°C, 30 s at 60°C, 3 min at 72°C) 14 times; 10 min at 72°C. Finally, nested PCR products were purified with 0.7X SPRI beads and subjected to Nanopore library preparation and sequencing.

For transcriptome analysis, 25% of 10x cDNA libraries were fragmented and processed for generating conventional gene expression library, according to the Single Cell 3’ Reagent Kit protocols (v3.1). Libraries were sequenced on an Illumina NovaSeq 6000 platform (paired-end 150 bp, PE150).

### Detection of DNA barcodes in bulk samples

Embryos were dissected at E18.5, and organs were harvested as below. Briefly, the placenta was divided into inner/outer sections at the labyrinth boundary. The brain was sectioned along the midline and cortical edge to delineate the anterior-posterior axis. In the liver, four lobes were collected separately as experimental replicates. Following removal of excess adipose tissue, the gut was divided into anterior, middle, and posterior segments. Tissues were washed in PBS to remove blood contaminants, flash-frozen in liquid nitrogen, and stored at -80°C.

Genomic DNA was extracted using proteinase K digestion and heat inactivation. Barcoding cassettes were amplified via PCR using primers #2426 and #2702, as previously described^45^. PCR primers incorporate 8-bp sample identifiers. PCR products were purified using 0.7x SPRI beads (Beckman Coulter, B23318), eluted in nuclease-free water, and quantified using Qubit™ 1X dsDNA High Sensitivity kit (Thermo, Q33231). Equimolar pools were prepared for Nanopore library preparation and sequencing (Oxford Nanopore Technologies).

### Single-cell multi-omic profiling of lineage barcodes, transcriptomes, and chromatin accessibility

In single-cell multi-omic experiments, *Rosa26^CreERT2/DeepTrack^* embryos were barcoded pre-gastrulation and harvested at E9.5. Single-cell suspensions were generated by tissue digestion with TrypLE Express and treated with 0.2-0.5x lysis buffer (10x Genomics Nuclei Isolation protocol, CG000365/CG000375) to isolate nuclei. Lysis progression was monitored via microscopy at 30-second intervals and halted with wash buffer to preserve nuclear integrity.

Lineage barcodes, transcriptomes, and chromatin accessibility were captured using the Chromium Next GEM Single Cell Multiome ATAC + Gene Expression Kit with the following modifications. A spike-in oligo (#ISPCR2999) was added to cDNA eluate prior to pre-amplification, and PCR program was adjusted to 3 min at 98°C; (15 s at 98°C, 20 s at 63°C, 3 min at 72°C) with 7 cycles; 3 minutes at 72°C. After cleanup, 35 µL of pre-amplified samples were used for cDNA amplification, 40 µL for ATAC library construction, and 31 µL for lineage barcode amplification. RNA and ATAC libraries were sequenced using Illumina NovaSeq 6000 platform (PE150). Lineage barcodes were amplified from cDNA using primer #ISPCR2999 (4 μL added) under the PCR condition as above. PCR products were purified using 0.7X SPRI beads and subjected to Nanopore sequencing.

### Generation of *Cdx2*-mutant tdTomato^+^ mouse embryonic stem (ES) cells

For *Cdx2* knock-out, plasmids encoding CMV-Cas9 (6 μg) and U6-*Cdx2*-sgRNA targeting *Cdx2* (3 μg) were co-electroporated into tdTomato^+^ ES cells using the Mouse Embryonic Stem Cell Nucleofector™ Kit (Lonza, VPH1001). Electroporation was performed in 100 µL mouse ES cell Nucleofector™ solution according to manufacturer protocols. Post-electroporation, cells were plated onto mouse feeder-coated 12-well plates. After 24 hours, ES cells were dissociated to single cells and replated at 500 cells per well in 6-well feeder-coated plates. Colonies were manually picked and expanded after 3-4 days of culture. Genotyped *Cdx2*-mutant and wild-type clones were retained to generate mutant and control chimeric mouse embryos.

### Mouse blastocyst collection, ES cell injection, and embryo transfer

Female ICR mice (4-6 weeks old) were superovulated via intraperitoneal injection of 5 IU pregnant mare serum gonadotropin, followed by 5 IU human chorionic gonadotropin (hCG) 48 hours later, and then crossed with male ICR mice. The females were checked for the vaginal plug in the following morning. Plugged females were sacrificed at E3.5 after hCG injection to collect blastocysts, which were cultured in KSOM (Millipore, MR-106-D) medium until ES cell injection.

*Cdx2* knockout ES cells described above were dissociated into single-cell suspension, and 10-12 cells were microinjected into each blastocyst. Injected blastocysts were briefly cultured in KSOM medium at 37°C and 5% CO_2_ for 1-2 hours. Fourteen to sixteen blastocysts were transferred into the uteri of 2.5 dpc pseudopregnant ICR mice (10-16 weeks). Chimeric embryos were analyzed at E10.5.

### Immunostaining

Mouse embryos were fixed overnight at 4°C in 4% paraformaldehyde (PFA, Solarbio, P1110). After washes with PBS, samples were permeabilized and blocked overnight at 4°C in PBS containing 0.5% Triton X-100 (v/v; Sigma, X100) and 10% fetal bovine serum (FBS). Primary antibodies (anti-T/Brachyury; CST, 81694S) were diluted 1:200 in permeabilization/blocking solution and incubated with embryos overnight at 4°C. Unbound antibodies were removed by washes with PBS containing 0.5% Triton X-100 at room temperature (RT). Secondary antibodies (Alexa Fluor 488-conjugated anti-rabbit; Jackson ImmunoResearch, 711-545-152) were diluted 1:500 in permeabilization/blocking solution and incubated samples for 1 hour at RT. After washes in PBS containing 0.5% Triton X-100, embryos were placed in PBS for imaging.

### Fluorescence-activated cell sorting (FACS)

Chimeric embryos (extraembryonic tissues removed) were harvested at E10.5, with somite counts between 23-25. Embryos were dissociated into single-cell suspensions via the lysis method described above and resuspended in PBS supplemented with 0.2% BSA. tdTomato^+^ and tdTomato^-^ populations were sorted using a BD FACS Fusion system and collected in PBS containing 0.04% BSA. Cell viability and concentration were assessed by acridine orange/propidium iodide (AO/PI) staining. Equal numbers of sorted cells were pooled for 10x Chromium scRNA-seq library preparation, followed by paired-end sequencing (PE150) on the Illumina NovaSeq 6000 platform.

### Bulk lineage barcode analysis

Nanopore sequencing was performed to resolve DeepTrack barcode sequences. Library construction and sequencing were conducted by Wuhan Benagen Technology Co., Ltd. Each read contains a lineage barcode cassette and a sample index for sample identification. To process reads, the sample index was extracted by aligning 5′ and 3′ anchor sequences flanking the index region in each read. Validated indices were matched against a whitelist to identify sample (organ) IDs. Reads were discarded if they exhibited >10% mismatch in anchor alignment or incorrect sample indices. Lineage barcodes were extracted using the RPBPBR pipeline^45^, and barcodes with only one read were excluded.

### Clonal identification via barcode generation probability (*Pgen*)

To enable high-resolution lineage tracing, we applied *Pgen* to identify clonal barcodes. *Pgen* denotes the probability that a specific barcode is generated through Cre-mediated recombination in the DeepTrack barcode cassette. This probability depends on the number of recombination steps required to form the barcode and the stochastic nature of recombination events. Lower *Pgen* values indicate rare barcodes that are less likely to arise independently in multiple cells. *Pgen* has been used for Polylox and PolyExpress barcoding systems in previous studies^22,33,45^. The threshold for *Pgen* was established using a binomial distribution model B (*k* | *N*, *p(b)*), where *N* is the total precursor cell count and *k* is the number of cells carrying a given barcode *b*. Given the limited number of cells (several hundred) at the labeling stage, a *Pgen* < 10^−3^ serves as a robust filter for single-cell-derived rare barcodes^12,45,47^.

### Fate classification of epiblast clones

Epiblast clonal fate bias was determined through the following steps. First, we merged the clonal barcode counts of tissues derived from the same germ layer, converting the clone-by-tissue matrix into a clone-by-germ-layer matrix. Next, clone-specific germ-layer contributions were normalized by the calculation of clone fraction in each germ layer using the R package FateMapper^73^. This generated a proportional matrix reflecting each clone’s bias toward different germ layers. Lastly, unsupervised K-means clustering (k=5) was applied to the normalized matrix, assigning epiblast clones into distinct fate-biased groups.

### Lineage tree reconstruction

Lineage trees were reconstructed using hierarchical clustering (McQuitty linkage) applied to lineage barcode similarity matrices. Cell type-cell type (single-cell lineage barcodes) similarities were computed with the ‘cospar.tl.fate_coupling’ function in the CoSpar Python package^49^.

### Single-cell RNA-seq data processing

scRNA-seq data were processed using CellRanger (v7.1.0; 10x Genomics). Mouse genome mm10 was set as the reference genome. Low-quality cells were filtered using the Scanpy Python package (v1.10.2)^74^, retaining cells with more than 200 detected genes, total UMI counts exceeding 1,000, and total UMI counts below the 95th percentile of the distribution. Cells with mitochondrial gene content exceeding 30% were removed. After quality control, the Scrublet Python package (v0.2.3) was applied with default parameters to identify and exclude potential doublets^75^. Batch correction and dimensional reduction were implemented via the scVI model from python package scvi-tools (v1.1.5), with the ‘n_latent’ parameter set to 10^76^. Unsupervised clustering analysis was performed using the ‘scanpy.tl.leiden’ function provided by Scanpy. We applied ‘scanpy.tl.umap’ function to visualize cellular clusters in two-dimensional space, with the ‘min_dist’ parameter set to 0.2^77^. Cell types were annotated using published marker genes^10,13^.

### Retrieval of single-cell lineage barcodes

Single-cell lineage barcodes were resolved from mRNA transcripts via Nanopore sequencing of the DeepTrack cassette integrated with cell indexes (CIs; 16 bp), introduced through indexed 10x Genomics beads during droplet encapsulation. Sequencing reads were first filtered to retain reads containing the expected structural elements: the 5’ and 3’ arms of the lineage barcode cassette, the cassette sequence itself, a poly-A tail, and the primer sequence to identify CIs. Reads lacking this validated architecture were discarded. Validated barcodes and CIs were computationally extracted using the RPBPBR pipeline^33,45^. After an additional filtering step to remove CIs absent from whole-transcriptome sequencing data and illegitimate lineage barcodes, remaining lineage barcodes were aggregated by CIs. In certain CI aggregates, individual cells contained multiple lineage barcodes, likely attributable to sequencing errors in CIs, PCR-generated chimeras, or doublets captured by the 10x Genomics. To assign a unique lineage barcode per cell, we used a filtering strategy based on read abundance, retaining only the highest-frequency barcodes for downstream analysis, as previously described^33^.

### Clonal fate classification of neuromesodermal progenitors (NMPs)

NMP clonal fate biases toward paraxial mesoderm (PXM) or spinal cord (SC) were quantified using FateMapper^73^, generating a normalized matrix reflecting the contribution of each NMP clonal barcode toward progeny cells. Unsupervised K-means clustering (k=3) was applied to the normalized matrix, assigning clones into bipotent NMPs, PXM-biased NMPs (PXM-NMPs), and SC-biased NMPs (SC-NMPs). Clonal barcodes exclusively detected in NMPs, with no progeny in differentiated lineages, were classified as self-renewing/undifferentiated.

### Spatial transcriptomic profiling and analysis

To spatially resolve transcriptomic identities across the mouse embryo, we applied Visium spatial transcriptomics (10x Genomics) to E9.5 embryos following manufacturer protocols (Visium Spatial Gene Expression User Guide, CG000239). Fresh embryos were flash-frozen in liquid nitrogen and embedded in optimal cutting temperature (OCT) compound. RNA quality of OCT-embedded embryo was assessed using an Agilent 2100 Bioanalyzer, and samples with RNA integrity number (RIN) ≥7 were selected for spatial transcriptomic profiling. A 10 µm frozen embryo section was mounted onto a Visium gene expression slide capture area. Tissue permeabilization (18 min) was followed by reverse transcription, cDNA library construction, and paired-end sequencing (Illumina NovaSeq 6000; PE150).

Raw sequencing data were processed using the Space Ranger pipeline (v.1.0.0; 10x Genomics) to align and summarize UMI counts according to the Mouse Genome mm10 reference. Gene expression matrices for spatial spots were normalized and scaled using the R package Seurat^78^. To infer the anterior-posterior axis distribution of spinal cord subpopulations, we integrated our scRNA-seq atlas with spatial transcriptomic data using the ‘run.RCTD’ function from the R package spacexr, with the parameter ‘doublet_mode’ set to ‘multi’^79^.

### NMP fate bias prediction by CoSpar

To predict fate outcomes for NMPs without detectable clonal barcodes, we integrated lineage barcodes with transcriptomes using CoSpar^49^. CoSpar leverages coherent, sparse of cell-state transitions to combine optimal transport theory with clonal data, enabling robust fate bias inference even when lineage information is incomplete. This approach inferred fate biases for NMPs lacking recovered barcodes by propagating clonal information across transcriptionally similar states. In brief, we defined NMPs as the early population and their progeny cells (paraxial mesoderm and spinal cord) as the late population. We then used ‘cospar.tmap.infer_Tmap_from_one_time_clones’ function to infer the transition probability matrix from NMPs to cells in the paraxial mesoderm and spinal cord. Based on the computed transition matrix, we applied ‘cospar.tl.fate_map’ function (default parameters) to infer the progenitor probability of each NMP differentiating to paraxial mesoderm or spinal cord. The progenitor probability, normalized by the maximum value among progenitors, indicated the likelihood of a given cell originating from a specific NMP. Pseudotime trajectories for NMP differentiation were reconstructed based on the transition matrix described above using the ‘cospar.pl.gene_expression_dynamics’ function.

### Identification of NMP gene modules

To identify high-confidence gene modules underlying cell fate determination, cells were aggregated into metacells (IReNA R package; ‘smoothByBin’ function), representing groups of cells with distinct transcriptional states^80^. Differentially expressed genes (DEGs) along pseudotime trajectories were identified using Monocle (v2.18.0; ‘differentialGeneTest’ function; adjusted *p* < 0.05; ≥10% metacell detection)^81^. DEGs were partitioned into five fate-associated modules via K-means clustering. Functional enrichment for each gene module was assessed using ClusterProfiler (v3.18.1; *p* < 0.05), based on the Gene Ontology (GO) database^82,83^.

### Single-cell ATAC-seq data processing

scATAC-seq data were processed using the R package ArchR (v1.0.1)^84^. In brief, ATAC fragment files were converted into Arrow files through ‘createArrowFiles’ function. Cells were filtered to ensure a minimum of 1000 fragments, a TSS enrichment score ≥ 4, and a blacklist ratio ≤ 0.05. Pseudo-bulk replicates were generated for each cell type, and peak calling was performed using ‘addReproduciblePeakSet’ function, with the parameter ‘cutoff’ set to 0.0001 and ‘peakMethod’ set to ‘MACS2’^85^. Transcription factor (TF) activity was computed using the ‘getVarDeviations’ function from ArchR, which quantifies changes in chromatin accessibility at TF binding sites to reflect TF regulatory activity^86^. Pseudobulk genomic tracks were visualized with the Integrative Genomics Viewer (IGV)^87^.

### Differential feature analysis between PXM-NMPs and SC-NMPs

DEGs and TF activities between PXM-NMPs and SC-NMPs were identified using ‘FindAllMarkers’ in Seurat^78^, with adjust *p* < 0.05 and absolute value of Log2FoldChange > 0.5. Differentially accessible regions (DARs) were identified by ‘getMarkerFeatures’ function from R package ArchR^84^, selecting DARs with *p* < 0.01 and Log2FoldChange > 2.

### Inference of fate-associated gene regulatory networks (GRNs)

GRNs governing NMP differentiation were reconstructed through the following analytical pipeline. PXM-NMPs and SC-NMPs were defined using CoSpar, as described above, in the single-cell multi-omic data. *Cis*-regulatory regions were mapped to nearby genes within 500 kb in NMPs and their progeny cells using the ‘LinkPeaks’ function from Signac^57^. To identify TF-target relationships in fate-divergent NMPs, TF binding motifs in *cis*-regulatory regions were identified via the ‘motif_analysis’ module in CellOracle based on ‘gimme.vertebrate.v5.0’ motif database^56,88^, and the strengths of TF-target regulation were inferred using the ‘get_links’ function in CellOracle. Finally, the top 2000 TF-target relationships, ranked by the above calculated weights, were selected as the core GRNs. GRNs were visualized using Cytoscape (v3.9.1)^89^.

## Data availability

The raw sequence data reported in this paper have been deposited in the Genome Sequence Archive in National Genomics Data Center, China National Center for Bioinformation / Beijing Institute of Genomics, Chinese Academy of Sciences, and will be publicly released upon acceptance of the manuscript^90,91^.

## Acknowledgements

We thank all the members of the Pei laboratory for discussions and support; the staff at the Laboratory Animal Resources Center, High-Performance Computing Center, Flow Cytometry Core, Genomic Core and Microscopy Core at Westlake University for support with techniques; and S. Wang, R. Sun, N. Shalin, for discussions and suggestions. We also thank H.R. Rodewald (DKFZ) for providing mice. This work was supported by National Key R&D Program of China (2022YFA1105700 to W.P., 2022YFA1302700 to Y.Z.), National Natural Science Foundation of China (32450525 and 82270123 to W.P., 32370710 to Y.Z., 82472668 to F.L.), Zhejiang Provincial Natural Science Foundation of China (LR25C070001 to W.P.), Pioneer and Leading Goose Key R&D Program of Zhejiang Province (2024SSYS0033 and 2024SSYS0034), and the Education Foundation of Westlake University.

## Author contributions

C.G. with W.P. conceived the DeepTrack approach. C.G. built the DeepTrack construct for mouse generation. X.M.W. generated *Cdx2* mutant mESCs and generated mouse chimeras. C.G, X.M.W., X.X.H., S.Z., and X.W.H. performed embryonic dissections. J.J. led the computational analyses, atlas construction, and development of the analytical framework; C.G., C.S., and M.Z. contributed to bioinformatic analyses. W.Y., F.S., X.W., H.Z., Q.D., F.L., D.H., X.L., G.P., and S.C provided intellectual input and revised the paper. W.P., D.P., and Y.Z. supervised the study and wrote the paper with input from C.G. and J.J., and W.P. conceived the study.

---

**Extended Data Fig. 1.**
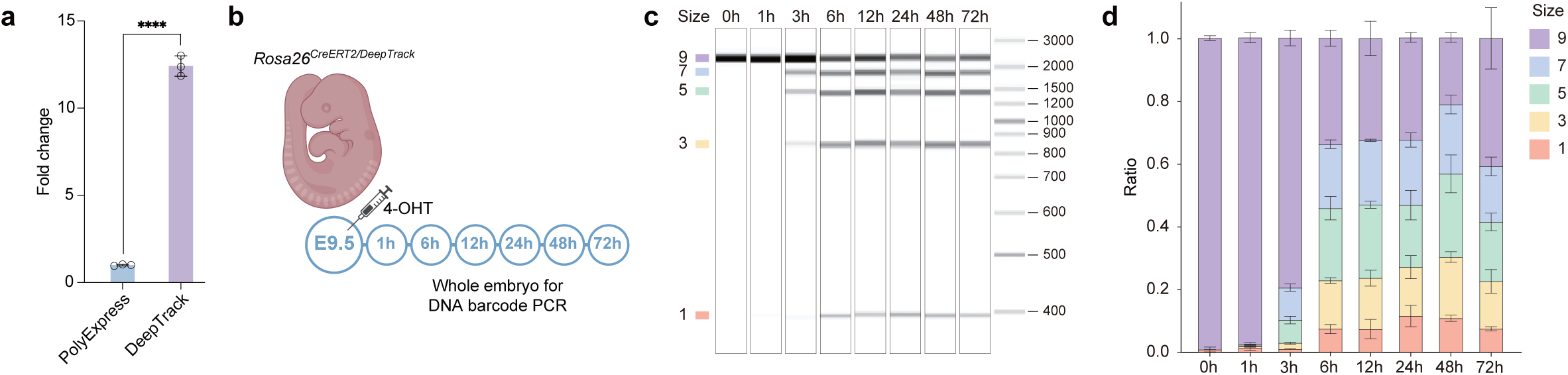
Characterization of DeepTrack. (**a**) Comparison of barcode expression in PolyloxExpress and DeepTrack mice (\*\*\*\**p* < 0.0001, *t* test). (**b**) *In vivo* barcoding in *Rosa26^CreERT2/DeepTrack^* mice for kinetic analysis. Barcodes were pulsed by 4-OHT administration to pregnant females at E9.5. Embryos were collected at indicated time points post induction (*n*=3 per time point). (**c**) Fragment analysis of barcode PCR products from embryos harvested at indicated post-induction time points. Barcode sizes are listed on the left. (**d**) Quantification of PCR band intensities corresponding to indicated barcode sizes.

**Extended Data Fig. 2.**
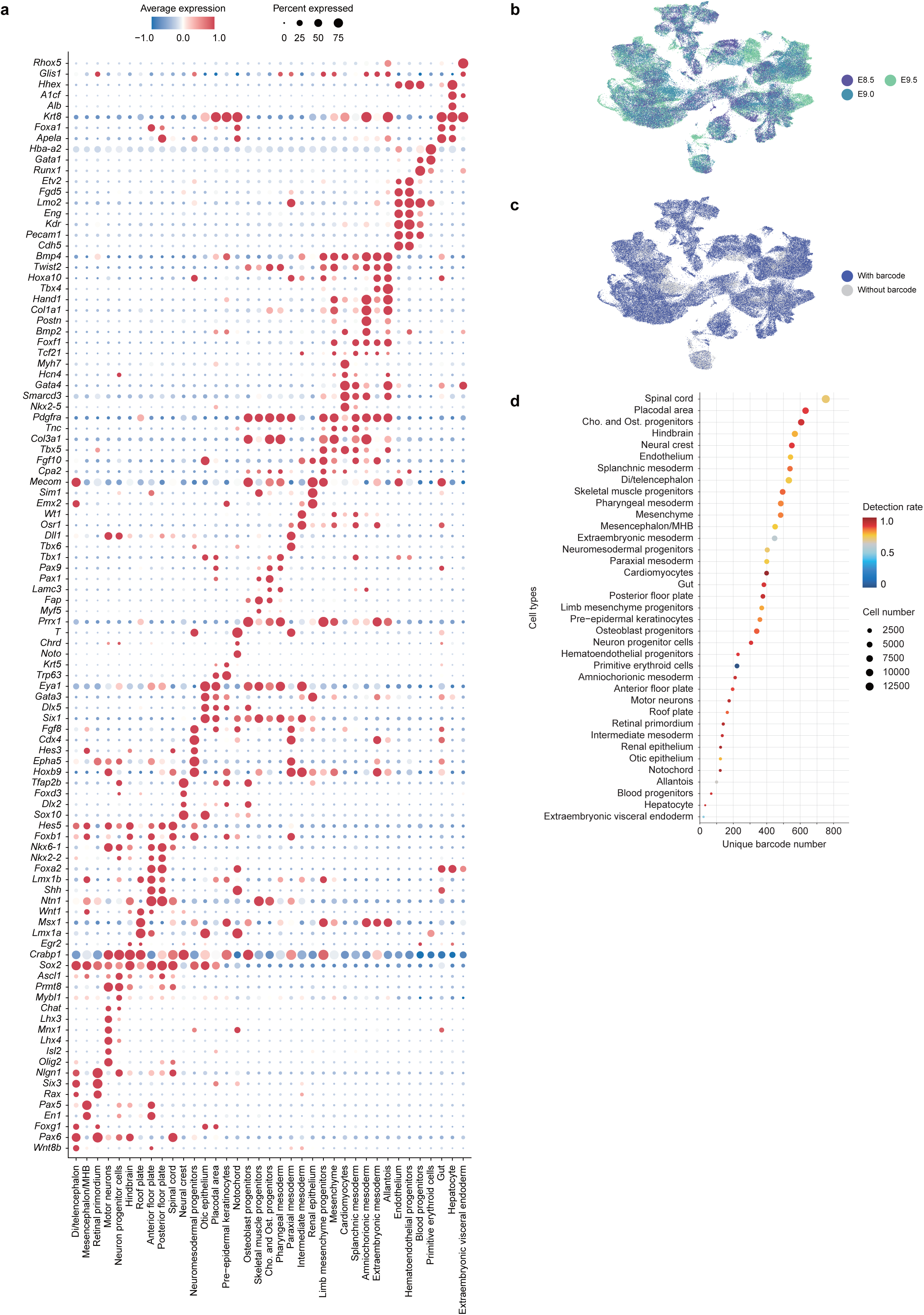
Characterization of single-cell clonal tracing during early organogenesis. (**a**) Marker genes used for cell type annotation in Fig. 1d. (**b**) UMAP displays cells from five embryos, color-coded by analyzed time points. (**c**) UMAP embedding highlights cells with lineage barcodes captured via scRNA-seq. (**d**) Unique barcode number, cell abundance (circle size), and barcode detection rate (color gradient) across annotated cell types.

**Extended Data Fig. 3.**
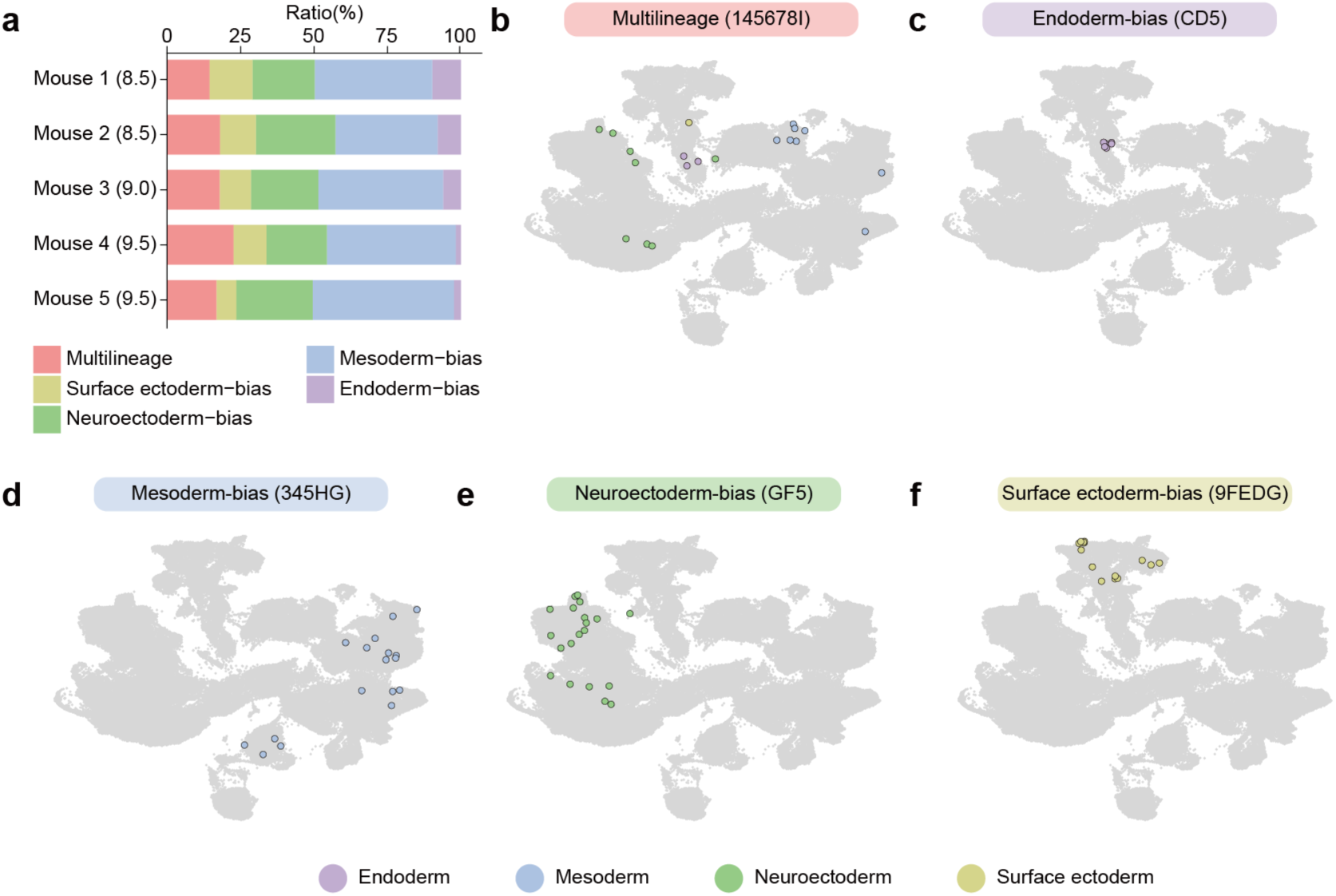
Clonal distribution during early organogenesis. (**a**) Overview of epiblast clonal fate patterns in individual embryos. (**b**-**f**) Representative clonal outcomes showing multilineage clones (**b**), and germ-layer biased clones in endoderm (**c**), mesoderm (**d**), neuroectoderm (**e**), and surface ectoderm (**f**).

**Extended Data Fig. 4.**
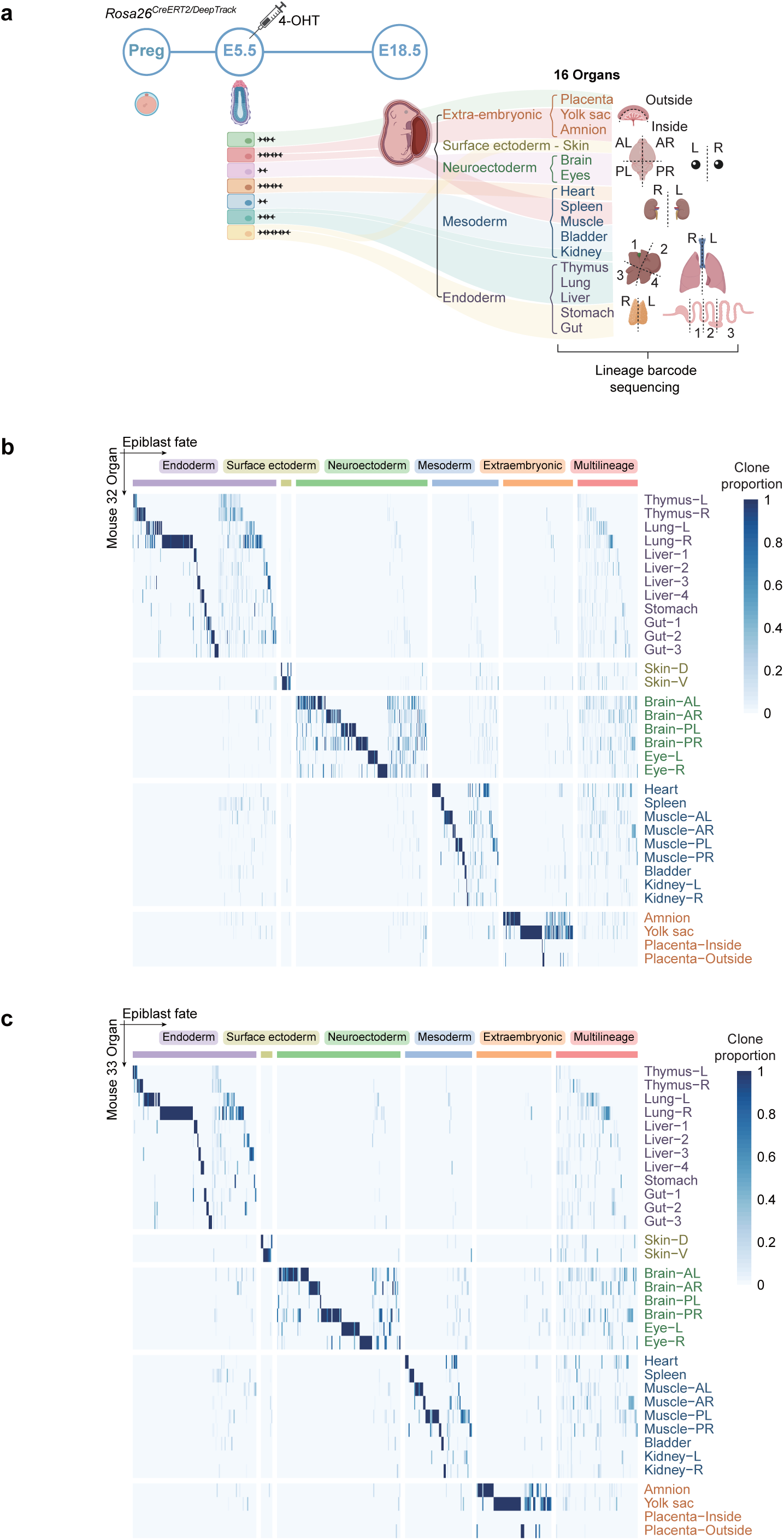
Bulk analysis of epiblast clonal contributions to fetal organs. (**a**) Lineage tracing workflow. Barcodes were induced in *Rosa26^CreERT2/DeepTrack^*embryos at E5.5 and analyzed in the indicated organs at E18.5. Numbers indicate sample replicates from the same organ. A, anterior; P, posterior; L, left; R, right; D, dorsal; V, ventral. (**b**-**c**) Heatmap showing fate outcomes of individual clones (columns, *Pgen* < 10^−3^) across tissues (rows) in two E18.5 embryos. Clonal fate biases (top boxes) were defined by K-means clustering, with color gradient reflecting clone proportion per tissue.

**Extended Data Fig. 5.**
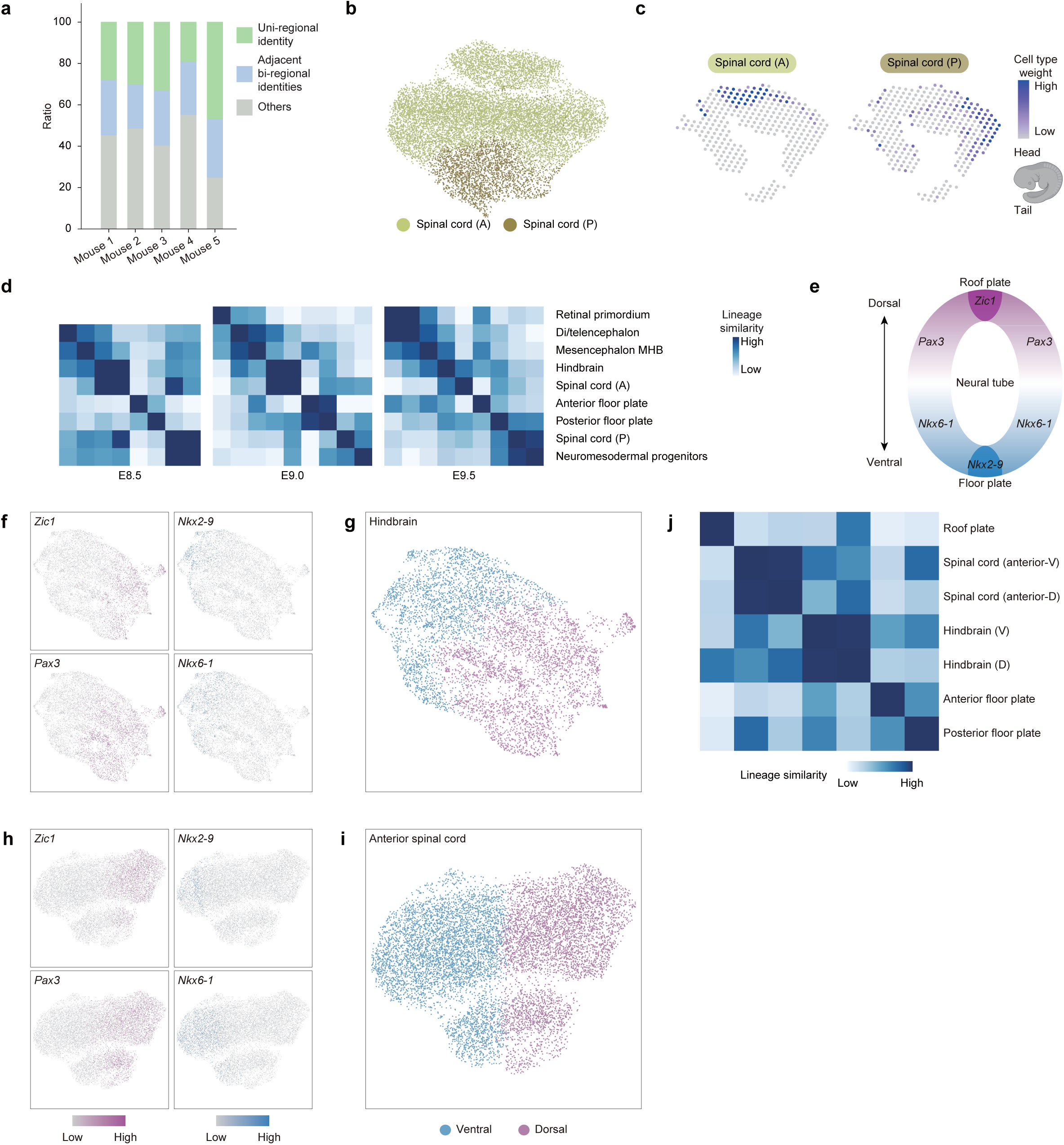
Spatiotemporal mapping of neuroectoderm development along the anterior-posterior and dorsal-ventral axes. (**a**) Clonal distribution across neural tube regions. Colored bars distinguish clones with uni-regional (green), adjacent bi-regional (blue), and other (gray) identities. (**b**) UMAP embedding of 12,597 spinal cord cells annotated with anterior (A) and posterior (P) identities. (**c**) Spatial distribution of anterior and posterior spinal cord subpopulations in an E9.5 embryo, with color gradient reflecting cell type weight across spatial spots. (**d**) Heatmap of lineage barcode similarity scores (color gradient) between nervous system regions at E8.5, E9.0, and E9.5. Single-cell data are aggregated as pseudobulk profiles. (**e**) Schematic illustrating spatial expression patterns of dorsal-ventral marker genes in E9.5 neural tube. (**f**-**g**) UMAP displays 6,041 hindbrain cells for showing dorsal-ventral patterning gene expression (**f**) and annotation of dorsal and ventral identities (**g**). (**h**-**i**) UMAP displays 10,136 anterior spinal cord cells for showing dorsal-ventral gene expression (**h**) and annotation of dorsal and ventral identities (**i**). (**j**) Heatmap of lineage barcode similarity scores across dorsal and ventral populations. Color gradient indicates similarity scores among cell types. D, dorsal; V, ventral.

**Extended Data Fig. 6.**
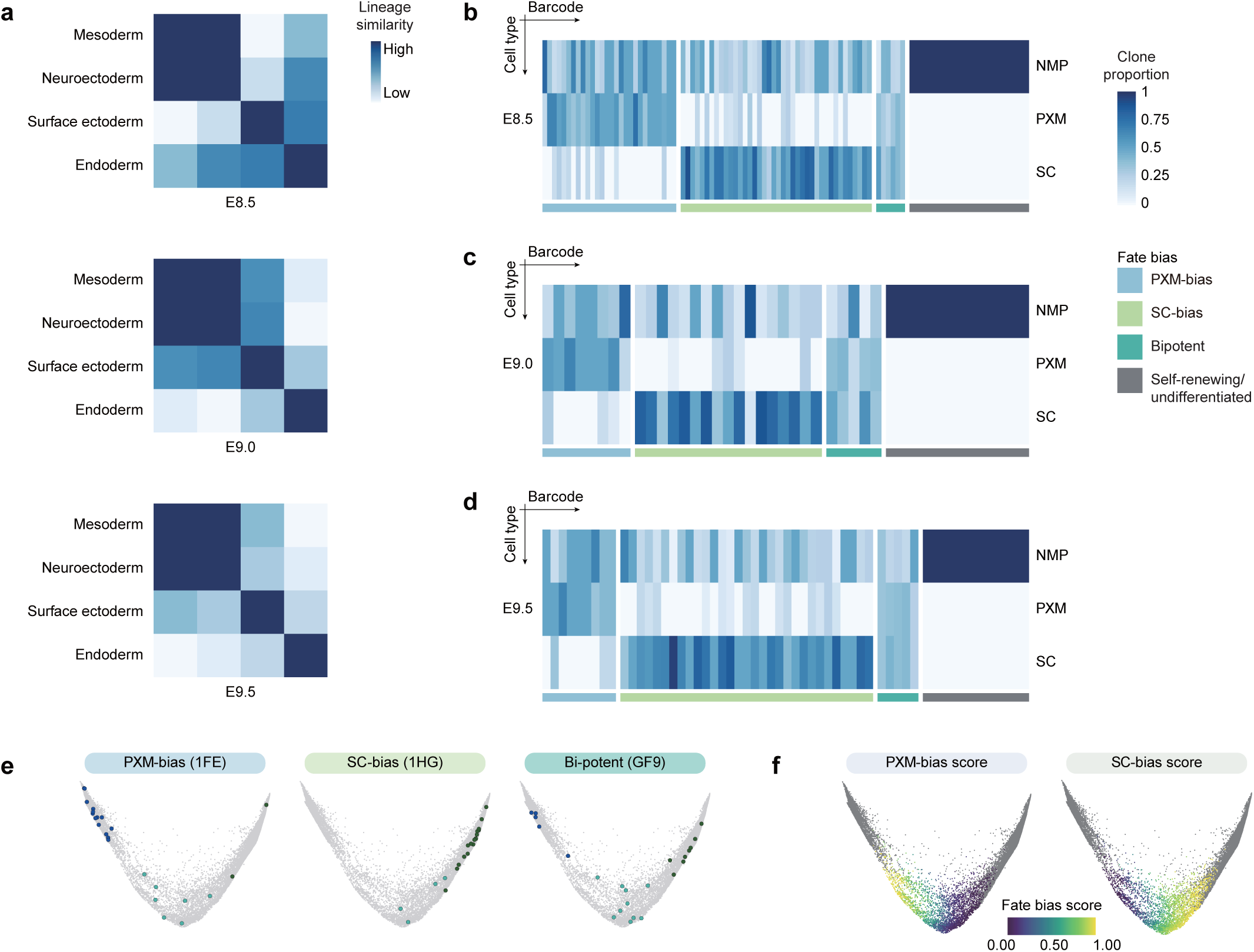
NMP clonal fate outcomes. (**a**) Heatmap of lineage barcode similarity among germ layer-derived cells at indicated time points, calculated from pseudobulk aggregation of single-cell data. Color gradient represents lineage similarity scores. (**b**-**d**) Distribution of clonal barcodes (columns) from NMPs to paraxial mesoderm (PXM) or spinal cord (SC) lineages (rows) at E8.5 (**b**), E9.0 (**c**), and E9.5 (**d**). Only the 200 barcodes labeling NMPs are displayed. Clonal fate biases (bottom boxes) were classified by K-means clustering, with color intensity reflecting clone proportion per cell type. (**e**) Representative clonal outcomes showing lineage-biased or bipotent clones. (**f**) CoSpar-predicted fate bias scores for NMPs transitioning to mesodermal (left) or neural (right) lineages.

**Extended Data Fig. 7.**
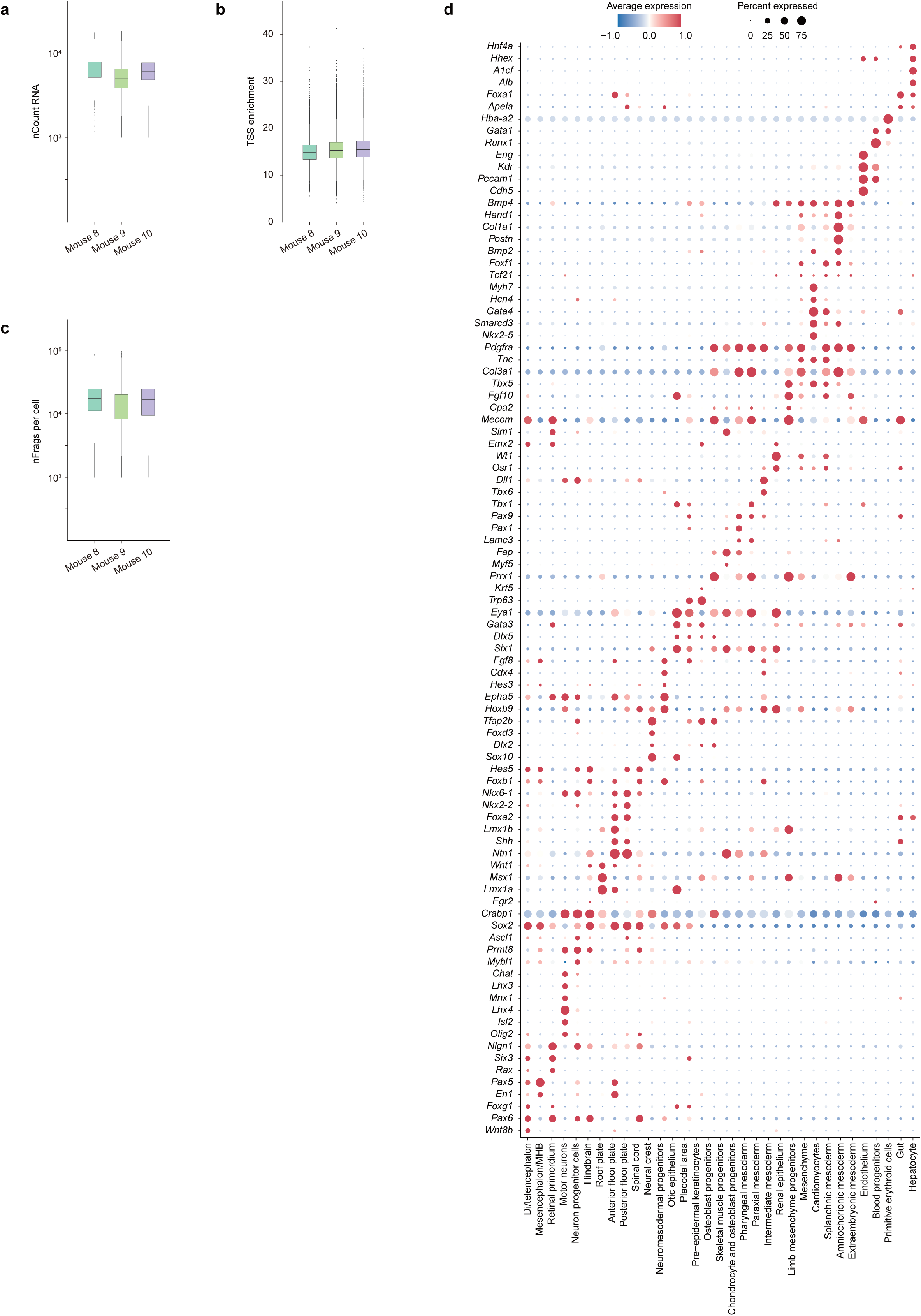
Data quality control and cell-type annotation for single-cell multi-omic clonal tracing datasets. (**a**-**c**) Various single-cell multi-omics quality metrics, split by biological replicates. (**d**) Marker genes used for cell type annotation in Fig. 3b.

**Extended Data Fig. 8.**
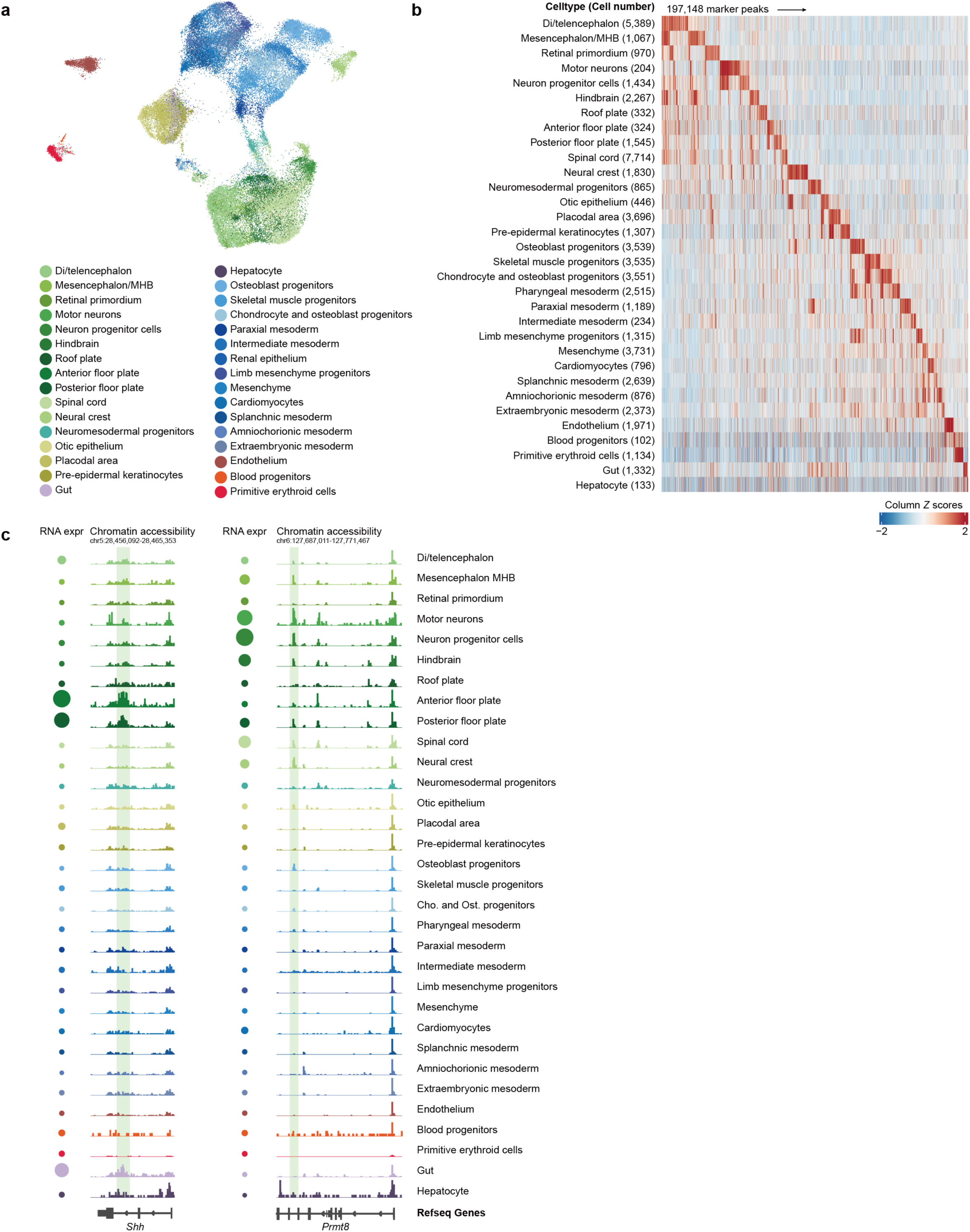
ATAC modality of single-cell multi-omic clonal tracing data. (**a**) UMAP of epigenome data from single-cell multi-omic clonal tracing datasets (60,355 cells), color-coded by annotated cell types. (**b**) Heatmap of scATAC-seq marker peaks (197,148) across embryonic cell types, with color scale representing column Z-scores of normalized accessibility. (**c**) RNA expression and chromatin accessibility of *Shh* (left) and *Prmt8* (right) across embryonic cell types. Intronic regions (highlighted by green bars) exhibit dynamic, cell-type specific accessibility patterns. Size of circle shows the gene expression level (left).

**Extended Data Fig. 9.**
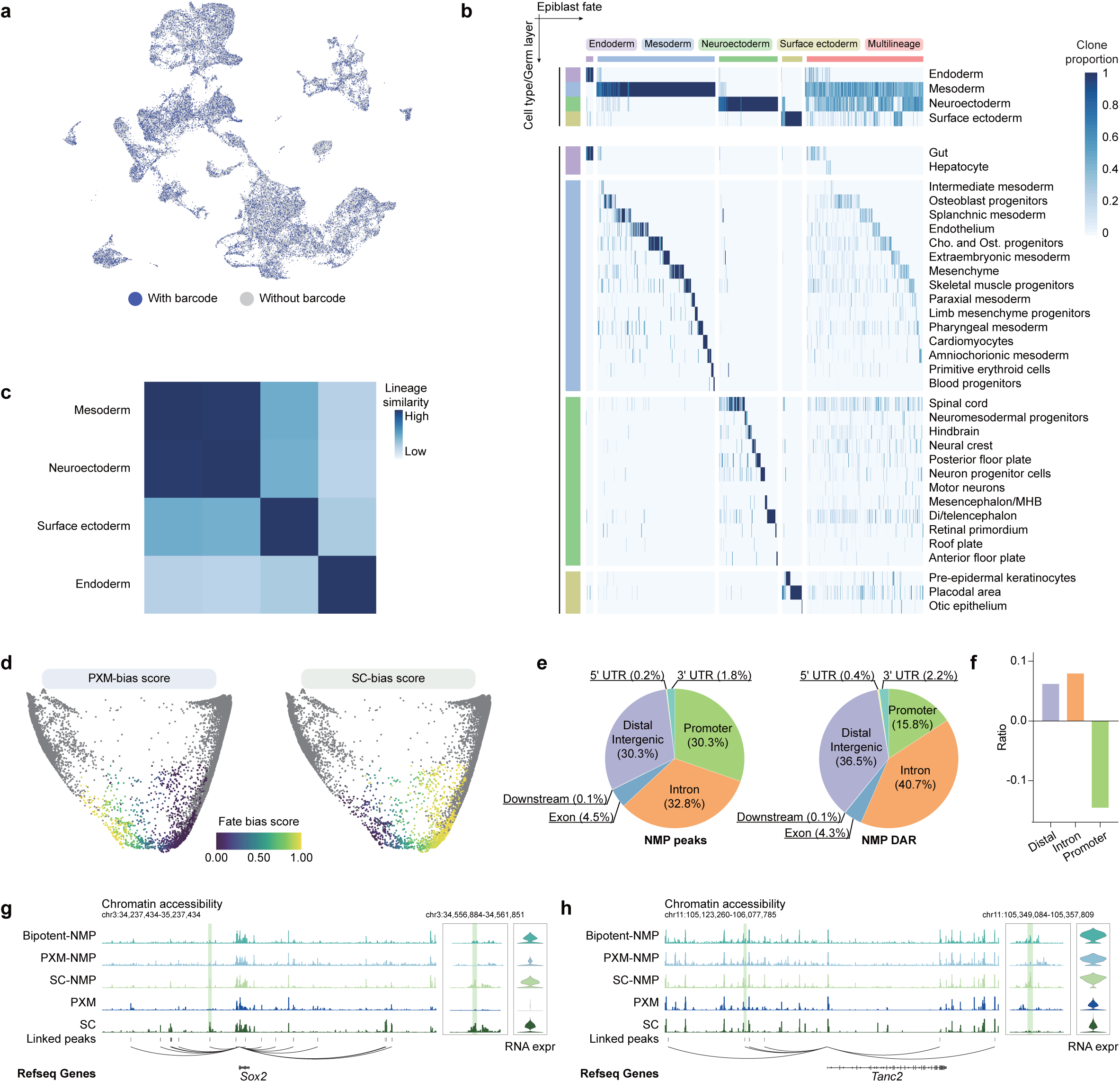
Clonal fate bias and fate-associated *cis*-regulatory elements in multi-omic lineage tracing data. (**a**) UMAP displays cells with lineage barcodes detected via single-cell multi-omic sequencing. Cells with barcodes (25,982 of 60,355 total cells) are highlighted in blue. (**b**) Clone distributions across germ layers and cell types in two embryos. Columns represent individual clones, and rows denote cell types. Clonal fate biases (top boxes) were defined by K-means clustering, with color intensity reflecting clone proportion per cell type. (**c**) Heatmap of lineage barcode similarity for distinct germ layers, calculated from pseudobulk aggregation of single-cell multi-omic data. Color gradient represents lineage similarity scores. (**d**) CoSpar-predicted fate bias scores for NMPs transitioning to mesodermal (left) or neural (right) lineages, distinguishing PXM-NMPs and SC-NMPs (1,561 cells). (**e**) Genomic distribution of accessible regions in NMPs (left) versus NMP fate-associated DARs (right). (**f**) Bar plots illustrating enrichment of fate-associated DARs in intronic and distal intergenic regions over promoters. (**g**-**h**) Chromatin accessibility and gene expression of *Sox2* and *Tanc2*. Tracks display pseudobulk ATAC-seq signals for three NMP subtypes and their progeny cells (PXM and SC). DARs enriched in SC-NMPs (green bars) are highlighted in the middle plot.

**Extended Data Fig. 10.**
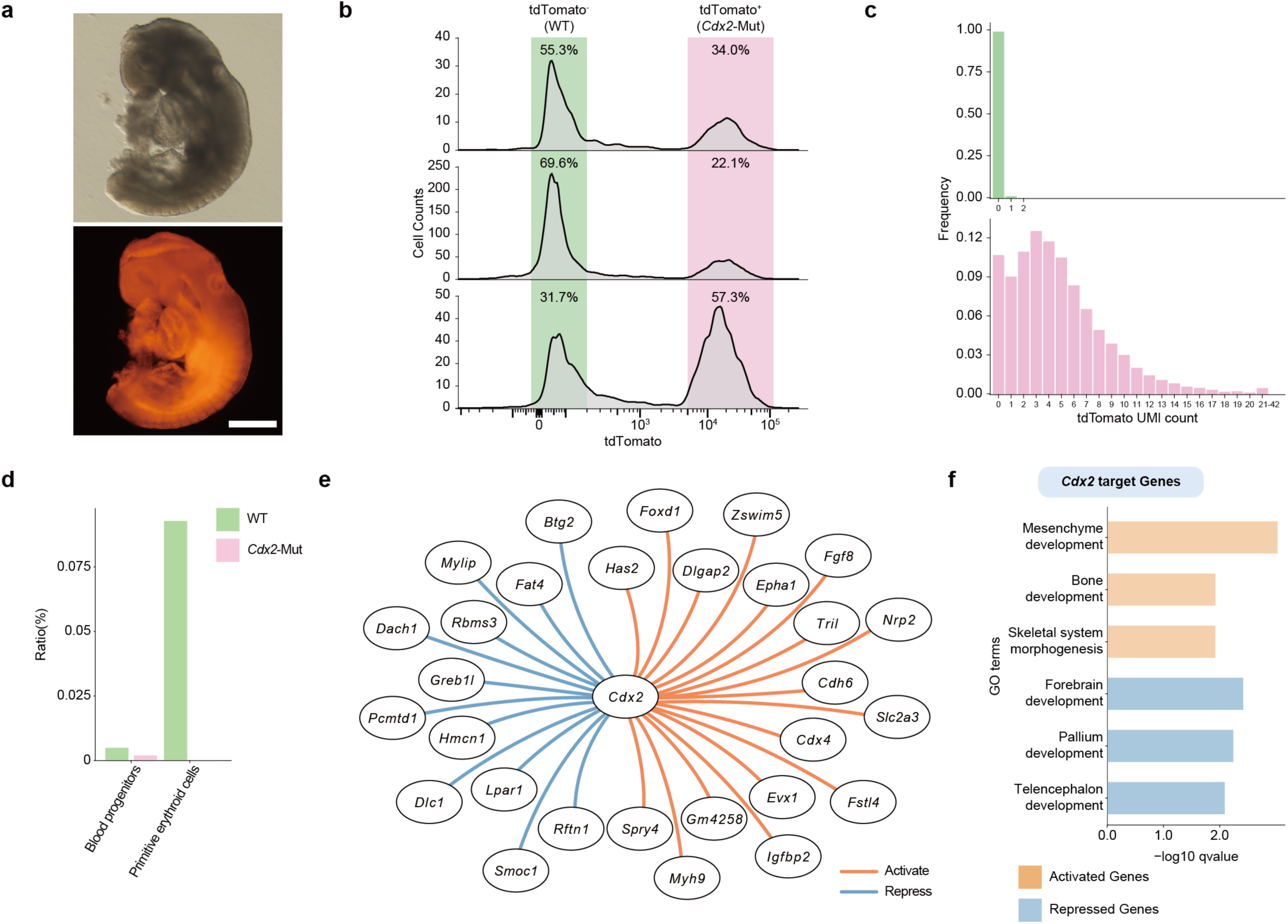
Characterization of *Cdx2*-mutant chimeric embryos. (**a**) Representative image of a chimeric embryo generated using *Cdx2*-mutant tdTomato^+^ embryonic stem cells. Scale bar, 500 μm. (**b**) Proportions of wild-type (WT; tdTomato^-^) and *Cdx2*-mutant cells (tdTomato^+^) quantified by flow cytometry analysis in three chimeras used for scRNA-seq (see Fig. 5a). (**c**) Histograms of tdTomato UMI counts in WT (top) and *Cdx2*-mutant cells (bottom) from scRNA-seq data. (**d**) Quantified contributions of wild-type and *Cdx2*-mutant cells to blood progenitors and primitive erythroid cells. (**e**) *Cdx2*-activated (red) and -repressed (blue) target genes within the fate-associated GRNs of PXM-NMPs. (**f**) GO enrichment analysis of *Cdx2*-activated and -repressed genes in the GRNs of PXM-NMPs.

